# Long-Term historical and Projected Herbivore Population Dynamics in Ngorongoro Crater, Tanzania

**DOI:** 10.1101/542910

**Authors:** Patricia D. Moehlman, Joseph O. Ogutu, Hans-Peter Piepho, Victor Runyoro, Mike Coughenour, Randall Boone

## Abstract

All the authors contributed to the analysis and interpretation of the data, drafting the article or revising it critically for important intellectual content. All authors read and approved the manuscript.

**Abstract:** The Ngorongoro Crater is an intact caldera with an area of approximately 310 km^2^. Long term records on herbivore populations, vegetation and rainfall made it possible to analyze historic and project future herbivore population dynamics. In 1974 there was a perturbation in that resident Maasai and their livestock were removed from the Crater. Vegetation structure changed in 1967 from predominately short grassland to mid and tall grasses dominating in 1995. Even with a change in grassland structure, total herbivore biomass remained relatively stable from 1963 to 2012, implying that the crater has a stable multi-herbivore community. However, in 1974, Maasai pastoralists were removed from the Ngorongoro Crater and there were significant changes in population trends for some herbivore species. Buffalo, elephant and ostrich numbers increased significantly during 1974-2012. The zebra population was stable from 1963 to 2012 whereas numbers of other eight species declined substantially between 1974 and 2012 relative to their peak numbers during 1974-1976. Numbers of Grant’s and Thomson’s gazelles, eland, kongoni, waterbuck (wet season only) declined significantly in the Crater in both seasons after 1974. Wildebeest numbers decreased in the Crater between 1974 and 2012 but this decrease was not statistically significant. In addition, some herbivore species were consistently more abundant inside the Crater during the wet than the dry season. This pattern was most evident for the large herbivore species requiring bulk forage, comprising buffalo, eland, and elephant. Analyses of rainfall indicated that there was a persistent annual cycle of 4.83 years. Herbivore population size was correlated with rainfall in both the wet and dry seasons. The relationships established between the time series of historic animal counts in the wet and dry seasons and lagged wet and dry season rainfall series were used to forecast the likely future trajectories of the wet and dry season population size for each species under three alternative climate change scenarios.

## Introduction

The Ngorongoro Crater, Tanzania is known world-wide for the abundance and diversity of its wildlife. It is situated in the Crater Highlands and is linked both to this area and the Serengeti Plains by the seasonal migration of several herbivores [1,2] and the emigration and immigration of large carnivores [4-8].

Since 1963, the herbivore population of Ngorongoro Crater has been monitored by the Ngorongoro Conservation Area Authority (NCAA), The College of African Wildlife Management and research scientists [1-3,9-12]. Since 1978, the Ngorongoro Ecological Monitoring Program has been responsible for conducting the wet and dry season censuses. The complete data set covers a period of 50 years (1963-2012). This data set makes it possible to assess long-term population trends and the stability of this multi-species wild herbivore community.

Earlier analyses indicated that the eviction of Maasai, the removal of their livestock and changes in rangeland management correlated with complex changes in vegetation composition and structure and wild herbivore populations. Previous papers have hypothesized that the removal of the Maasai pastoralists was a key factor in changes observed in herbivore populations. Pastoral pasture management may have affected vegetation structure and species composition [12,13].

This paper further examines the hypothesis that the removal of the Maasai and their livestock from the Crater in 1974 affected the plant structure in the crater and the population dynamics of the resident wild herbivore species depending on their life-history traits (body size, gut morphology) and life-history strategies (feeding style, foraging style, and movement patterns).

In addition we examine the hypothesis that rainfall variation influences the herbivore population dynamics and density, differentiated by life-history traits and strategies. Extreme rainfall in the Crater, which waterlogs large parts of the Crater should adversely affect wildlife, just like droughts, if large parts of the Crater become waterlogged. Additionally, high rainfall promotes excessive grass growth and dilutes plant nutrients, hence reducing vegetation quality for herbivores.

Relationships established between historic population abundance and historic rainfall are used to project the impacts of three different future rainfall scenarios on wild herbivore population dynamics to 2100.

## Methods

### Study Area

Ngorongoro Crater, Tanzania is known world-wide for the abundance and diversity of its wildlife. The crater (3°10′ S, 35° 35′ E) is a large intact caldera with an area of approximately 310 km^2^. The floor of the crater is about 250 km^2^ (1,700 m above sea level) and the sides rise steeply 500 meters to the rim. The geology, soils and vegetation of the crater were described by Herlocker and Dirschl [14] and Anderson and Herlocker [15]. The crater has the largest catchment basin in the Ngorongoro Highlands [16] and receives water from Lalratati and Edeani streams and Lerai spring from Oldeani Mountain to the south. Seneto spring provides water to Seneto swamp and Lake Magadi from the southwest. Olmoti Crater provides runoff to Laawanay and Lemunga rivers in the north, which supply Mandusi swamp and Lake Magadi. Lljoro Nyuki river, in the northeast provides water to Gorigor swamp. Ngaitokitok spring in the eastern part of the crater also supplies Gorigor swamp and Lake Magadi. Soil characteristics and drainage affect vegetation species and during the dry season soil moisture is dependent on the crater’s catchment system (Fig. 1). The wildlife of Ngorongoro Crater has had a protected status since 1921. In 1974, resident Maasai pastoralists, their bomas and livestock were removed from the crater [10,17]. The area has been administered by the Ngorongoro Conservation Unit since 1959 and by the Ngorongoro Conservation Area Authority since 1975 as part of a protected multiple land use area (8,292 km^2^).

**Fig 1.**
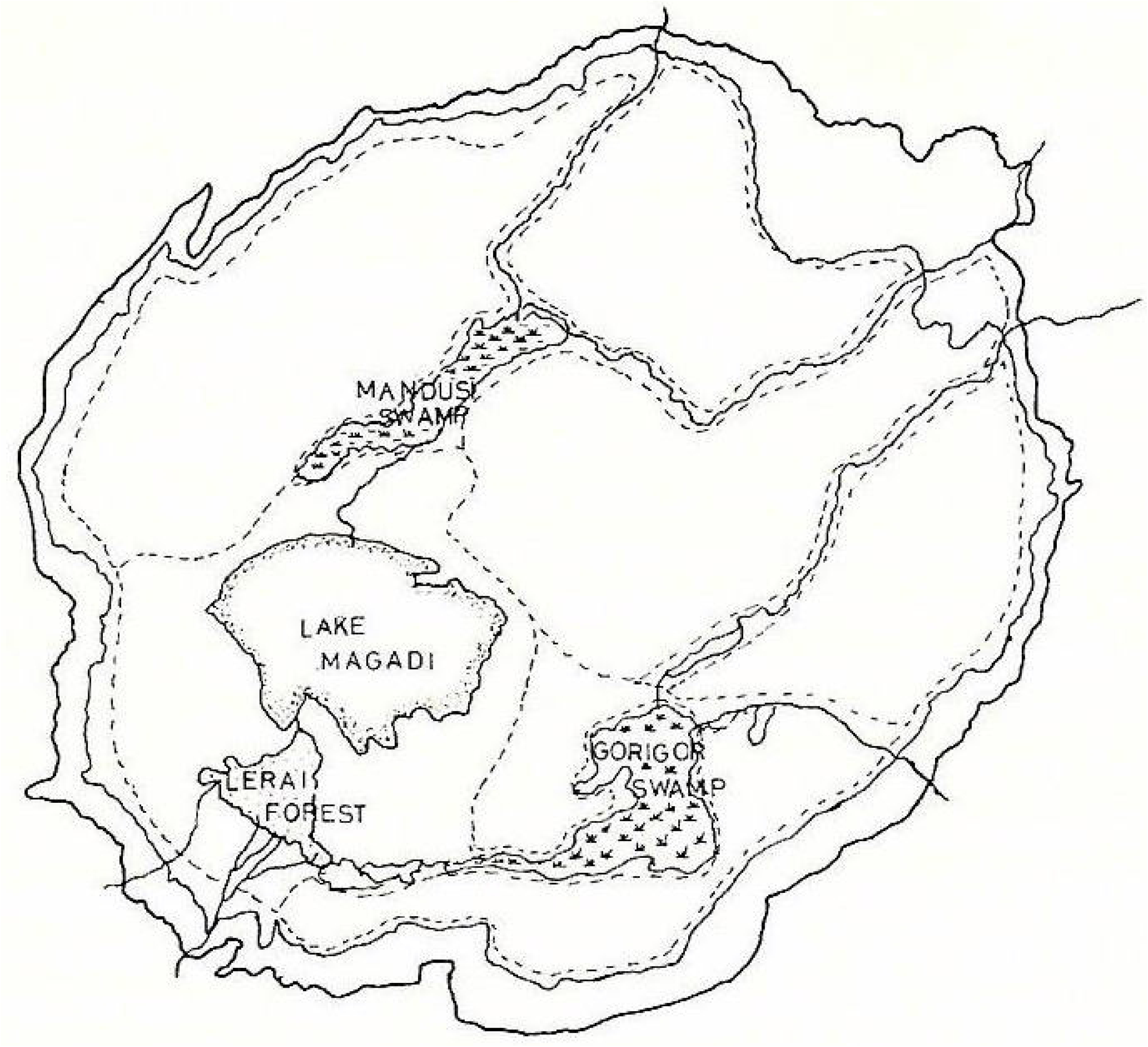
Ngorongoro Crater and Census Blocks [12]).

### Wild herbivore populations

Long term data sets were available for eleven mammalian herbivores, i.e. Wildebeest (*Connochaetes taurinus*), Plains Zebra (*Equus quagga)*, Cape Buffalo (*Syncerus caffer*), Thomson’s gazelle (*Eudorcas thomsonii*), Grant’s gazelle (*Nanger granti*), Eland (*Tragelapus oryx*), Kongoni (*Alcelaphus buselaphus*), Waterbuck (*Kobus ellipsiprymnus*), Warthog (*Phacochoerus aethiopicus)*, Elephant (*Loxodonta africana*) and Black Rhino (*Diceros bicornis*) and one bird, the Ostrich (*Struthio camelus*).

Zebra are mid-sized herbivores, but they are non-ruminants. Hence they are not limited by a four-chambered stomach system and can opt to consume larger amounts of higher fiber (lower quality) grasses to meet their nutritional requirements [18].

Buffalo are large bodied ruminants and although they require a larger amount of food per individual, the quality can be lower and they can tolerate a higher proportion of fiber in their diet [18,19]. Buffalo prefer longer grass and select for a high ratio of leaf to stem [20].

The herbivores were classified into functional categories, i.e., grazers (Thomson’s gazelle, kongoni, wildebeest, eland, buffalo, and zebra) and mixed browsers/grazers (Grant’s gazelle, waterbuck, black rhino and elephant) [21-27]. The ostrich is primarily a herbivore, but will also eat invertebrates and occasionally rodents [28].

### Herbivore Total Counts

Since 1963, the herbivore population of Ngorongoro Crater has been monitored by the Ngorongoro Conservation Area Authority (NCAA), The College of African Wildlife Management and research scientists [1-3,9-12]. Since 1987, the Ngorongoro Ecological Monitoring Program has been responsible for conducting the wet and dry season censuses. This data set makes it possible to assess long-term population trends and the stability of this multi-species wild herbivore community. Here, we consider the data set covering a period of 50 years (1963-2012).

Total counts of large mammals in the wet and dry seasons have been done in the crater since the 1963. The floor of the crater was divided into six blocks (Fig 1) that cover the entire area except for inaccessible areas, i.e., Lake Magadi, Lerai Forest and the Mandusi and Gorigor swamps. The ground censuses are done by one team per block composed of one driver, one observer and one recorder in a four-wheel drive vehicle driving along line transects that are one kilometer apart. Since 1987 each of the six teams has been supplied with a 1:50,000 map marked with the transects, a compass, binoculars and a mechanical counter. Each block takes six to eight hours to complete and all blocks are censused simultaneously [12, 29-34]. Unpublished records of NCAA and NEMP 1963 - 2012 provide most of the seasonal data on animal numbers.

From 1981 to 1985 there were no censuses. In 1986 the total counts were resumed and strip counts were used for counting gazelles and warthog and analyzed with Jolly’s method 2 [35]. Strip counts were discontinued after 1989 because of unacceptably large confidence limits and the difficulty of maintaining absolutely straight transects in the wet season. Data from strip counts were not used in the analyses. However, ‘transects’ are still used to ensure complete coverage of each block. Total aerial counts were conducted in 1964, 1965, 1966, 1977, 1978 and 1988 [12]. A systematic reconnaissance flight count was done in 1980 which included warthogs for the first time [36]. Total counts of warthogs started in 1986 [12]. The count totals for the 12 most common large herbivore species for the Ngorongoro Crater during 1963-2012 are provided in S1 Data. The same data set with the missing counts imputed using a state space model is provided in S2 Data.

Total biomass for the wet and dry seasons for each year were calculated using unit weights in Coe et al. [37]. Biomass was calculated separately for each species and season. The fact that black rhinos, elephants and warthogs move into the forest at the edge of the crater and into Lerai forest make them more difficult to count and may affect their contribution to biomass.

### Vegetation

Changes in vegetation composition and structure were measured by digitizing and comparing vegetation maps that were done in 1966-67 and 1995 [14,38]. Maps were digitized in ArcGIS 9.1 (ESRI, Redlands, California) and projected to UTM Zone 36, WGS 1984 datum. Attributes on the maps were digitized, and in both maps the plant height for primary and secondary canopy species was used to determine the presence of short, mid, mid-tall and tall grass structure.

Fires were suppressed from 1974, when the Maasai were removed, until 2001 [39]. Prescribed burning started in 2001. Transects were used to measure canopy height and biomass in kg/ha estimated by linear regression. Starting in 2001, areas with more than 4000 kg/ha were burned every year at the end of the dry season (September/October). It was recommended that 10-20% of the crater floor was burned on a rotational basis. Highest tick density occurred in the peak dry season (September/October) in the longest grass. Twenty-seven months after the start of prescribed burning, there was a significant decrease in tick density in burned areas. Short grass (<10 cm) areas with a fuel load of less than 4000 kg/ha appear to correlate with limited tick survival [39]. From 2002 to 2011 there was prescribed burning but no records were maintained. From 2012 to 2017 approximately 10 to 15 km^2^ were burned each year in different areas. In 2012-2015 burning was done in the northern and northeastern portion of the crater. In 2016 and 2017 burning was conducted in the eastern and then the east central portion of the crater (Pers comm NCAA 2018).

### Rainfall

Long-term rainfall data was not available for the crater floor. We therefore used monthly rainfall measured from 1964 to 2014 at Ngorongoro Headquarters on the southern rim of the crater. The rainfall recorded at the Ngorongoro Conservation Area Authority (NCAA) headquarters during 1963-2014 is provided in S3 Data.

### Projection of rainfall and temperature

Total monthly rainfall and average monthly minimum and maximum temperatures for Ngorongoro Crater were projected over the period 2013-2100 based on regional downscaled climate model data sets from the Coordinated Regional Climate Downscaling Experiment (CORDEX). Downscaling is done using multiple regional climate models as well as statistical downscaling techniques. Three climate scenarios defined in terms of Representative Concentration Pathways (RCPs) were used to project rainfall and temperatures for the Ngorongoro Crater. The three RCPs are RCP2.6, RCP4.5 and RCP8.5 in which the numeric suffixes denote radiative forcings (global energy imbalances), measured in watts/m^2^, by the year 2100. The RCP2.6 emission pathway (best case scenario) is representative for scenarios leading to very low greenhouse gas concentration levels [40]. RCP4.5 (intermediate scenario) is a stabilization scenario for which the total radiative forcing is stabilized before 2100 by employment of a range of technologies and strategies for reducing greenhouse gas emissions [41]. RCP8.5 (worst case scenario) is characterized by increasing greenhouse gas emission over time representative for scenarios leading to high greenhouse gas concentration levels [42]. Rainfall, minimum and maximum temperature projections were made for a 50 × 50 km box defined by longitudes (34.97, 35.7) and latitudes (−3.38, −2.787).

### Ethics Statement

All the animal counts in the Ngorongoro Crater were carried out as part of a long-term monitoring Program under the auspices of the Ngorongoro Conservation Area Authority (NCAA).

### Statistical modeling and analysis

#### Modeling trends in animal population size and biomass

Time trends in count totals for all the 12 most common large herbivore species were modeled simultaneously using a multivariate semiparametric generalized linear mixed model assuming a negative binomial error distribution and a log-link function. The variance of the negative binomial distribution model var(y) was specified as a quadratic function of the mean (μ), var(y) = μ(1 + μ/k), where k is the scale parameter. The semi-parametric model is highly flexible and able to accommodate irregularly spaced, non-normal and overdispersed count data with many zeroes or missing values. The parametric part of the model contains only the main effect of animal species to allow direct estimation of the average population sizes for the different species in each season. The non-parametric part of the model contains two continuous random effects, each of which specifies a penalized spline variance-covariance structure. The first random spline effect fits a penalized cubic B-spline (P-spline, [43] with a third-order difference penalty to random spline coefficients common to all the 12 species and therefore models the temporal trend shared by all the species. The second random spline effect fits a penalized cubic B-spline with random spline coefficients specific to each species and thus models the temporal trend unique to each species. Each random spline effect had 20 equally spaced interior knots placed on the running date of the surveys (1963,…, 2012) plus three evenly spaced exterior knots placed at both the start date (1963) and end date (2012) of the surveys. De Boor [44] describes the precise computational and mathematical properties of B-splines. The specific smoothers we used derive from the automatic smoothers described in Ruppert, Wand and Carrol [45].

The full model contains three variance components to be estimated, corresponding to the random spline time trend common to all species, random spline effects for the time trend specific to each species and the scale parameter for the negative binomial distribution. The full trend model was fitted by the residual penalized quasi-likelihood (pseudo-likelihood) method [46] in the SAS GLIMMIX procedure [47]. More elaborate details on this approach to modelling animal population trends can be found in Ogutu et al. [48]. Separate trend models were fit to the wet and dry season count totals for simplicity. The denominator degrees of freedom for Wald-type F-tests were approximated using the method of Kenward and Roger [49]. Temporal trends in total biomass calculated using unit weights in Coe et al. [37] were similarly modeled, separately for each season.

We used constructed spline effects to estimate and contrast population sizes for each species between 1964 versus 1974 when the Maasai and their livestock were evicted from the Crater and 1974 versus 2012. The constructed spline effects consisted of a cubic B-spline basis with three equally spaced interior knots. A constructed regression spline effect expands the original time series of animal survey dates into a larger number of new variables (seven in this specific case). Each of the new variables is a univariate spline transformation. The constructed spline effects are special model effects, in contrast to classical classification or continuous effects, and can be constructed using various other basis functions, including the truncated power function basis. These special model effects allowed estimation of the expected counts of each animal species at specified values of time (1964, 1974 and 2012). Because of the two comparisons made for each species, a multiplicity correction was made to control the familywise Type I error rate. We thus computed simulation-based step-down-adjusted *p*-values [50].

#### Modeling temporal variation in rainfall

The time series of rainfall was analyzed by using the unobserved components model (UCM), which is a special case of the linear Gaussian state space or structural time series model, to decompose the annual, wet season and dry season rainfall time series (*r*_*t*_) into their trend (*µ*_*t*_), cyclical (*φ*_*t*_), seasonal (*δ*_*t*_) and irregular (*ϵ*_*t*_) components

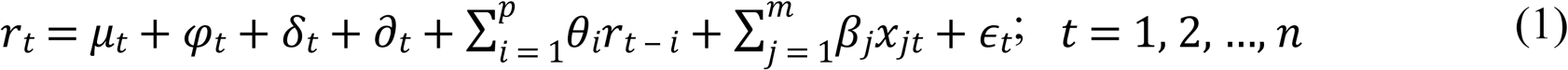

in which ∂_*t*_ is the autoregressive component,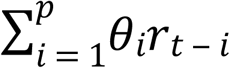 is the autoregressive regression terms, β_j_ are the explanatory regression coefficients, x_jt_ are regression variables treated as fixed effects and (*ϵ*_*t*_) are independent and identically (*i*.*i*.*d*.) normally distributed errors or disturbances having zero mean and variance 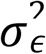. This is equivalent to assuming that *ϵ*_*t*_ is a Gaussian white noise process. The different model components are assumed to be statistically independent of each other.

We first assume a random walk (RW) model for the time trend, or equivalently that the trend (μ_t_) remains approximately constant through time. The RW trend model can be specified as

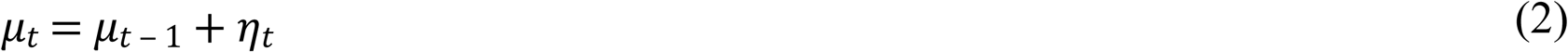

where 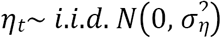. Note that 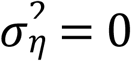 implies that *µ*_*t*_ = *a constant*.

Additionally, we assume a stochastic cycle (*φ*_*t*_) with a fixed period (*p* > 2), a damping factor (*ρ*) and a time-varying amplitude and phase given by

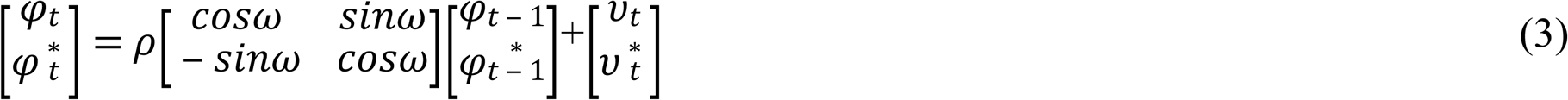

where 0 < *ρ* ≤ 1, *ω* = 2 × *π*/*p* is the angular frequency of the cycle, *υ*_*t*_ and 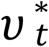 are independent Gaussian disturbances with zero mean and variance 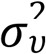 and 0 < *ω* < *π*. Values of *ρ, p* and 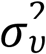 are estimated from the data alongside the other model parameters. The damping factor *ρ* governs the stationarity properties of the random sequence *φ*_*t*_ such that *φ*_*t*_ has a stationary distribution with mean zero and variance 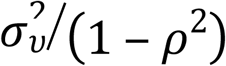 if *ρ* < 1 but is nonstationary if *ρ* = 1. We specified and tested for significance of up to three cycles in the annual, wet season and dry season rainfall components.

Besides the random walk model (2), we modelled the trend component using a locally linear time trend incorporating the level and slope components and specified by

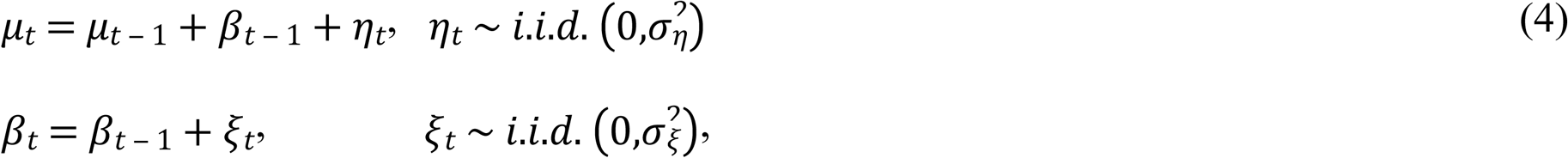

where the disturbance variances 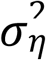 and 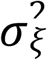 are assumed to be independent. The UCM models (1) and (4), without the seasonal and regression components, were fitted by the diffuse Kalman filtering and smoothing algorithm [51] in the SAS UCM procedure [47].

We grouped years with the annual rainfall falling within the 0–10, 11–25, 26–40, 41–75, 76–90, 91–95 and 96–100^th^ percentiles of the frequency distribution of the annual rainfall as extreme, severe or moderate drought years, normal, wet, very wet or extremely wet years, respectively. The dry (June to October) and wet (November to May) seasons were similarly grouped [52]. These percentiles allowed us to quantify the degree of rainfall deficit or surfeit and represent the expected broad transitions in rainfall influences on vegetation production and quality in each year and season.

#### Relating animal population size to rainfall

Rainfall primarily governs vegetation production and quality in savannas [53-55], and therefore also the aggregate and species-specific biomass levels of large African savanna ungulates [37,56-58]. Population size was related to moving averages of the annual, wet season and dry season rainfall components each computed over 1, 2,…, 6 years for a total of six different moving averages per rainfall component. The maximum of 6-year window was chosen to match the approximately 5-year dominant periodicity or quasi-cyclical pattern estimated for the time series of the wet season and annual rainfall components (Fig S3), based on the UCM model and spectral functions evaluated by the finite Fourier transform method. Spectral densities were obtained by smoothing the raw spectra or periodograms using moving average smoothing with weights derived from the Parzen kernel [47].

The moving rainfall averages index changing habitat suitability for ungulates associated with carry-over effects of prior rainfall on vegetation conditions. Population sizes was related to each of the 18 moving averages using a generalized linear model assuming a negative binomial error distribution and a log link function. The following six different functional forms were used for each of the 18 moving averages [58]:

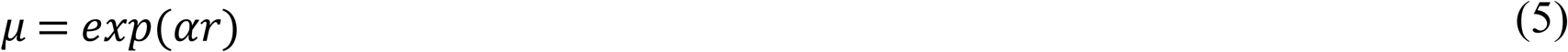

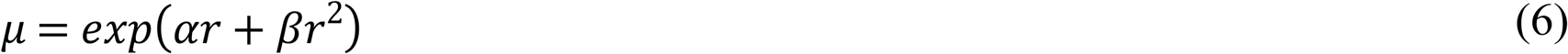

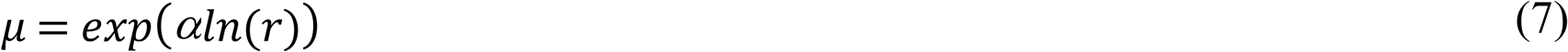

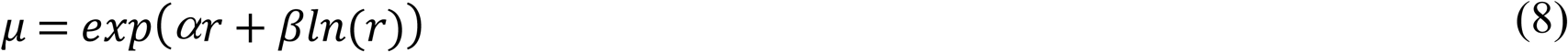

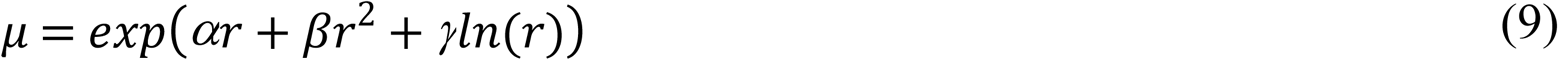

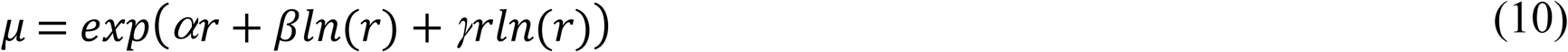

These models were selected to represent (1) a linear increase or decrease in animal population size with increasing rainfall, (2) an increase in animal abundance with increasing rainfall up to some asymptote, or (3) an increase in animal abundance with increasing rainfall up to a peak at some intermediate levels of rainfall, followed by decline with further increase in rainfall [58]. The most strongly supported rainfall component, specific moving average and functional form were then selected using the corrected Akaike Information Criterion (AICc, [59] Tables S13-S14).

#### Forecasting animal population dynamics using projected future climate

The relationships established between the time series of historic animal counts in the wet and dry seasons and lagged wet and dry season rainfall series were used to forecast the likely future trajectories of the wet and dry season population size for each species under three alternative climate change scenarios. We used the (Vector Autoregressive Moving Average Processes) VARMAX model to model the dynamic relationships between the wet and dry season counts of each species and the lagged wet and dry season rainfall and to forecast the seasonal animal counts. The model is very general and highly flexible and allows for the following among other features. 1) Modelling several time series of animal counts simultaneously. 2) Accounting for relationships among the individual animal count component series with current and past values of the other series. 3) Feedback and cross-correlated explanatory series. 4) Cointegration of the component animal series to achieve stationarity. 5) Seasonality in the animal count series. 6) Autoregressive errors. 7) Moving average errors. 8) Mixed autoregressive and moving average errors. 9) Distributed lags in the explanatory variable series. 10). Unequal or heteroscedastic covariances for the residuals.

The VARMAX model incorporating an autoregressive process of order p, moving average process of order q and in which the number of lags of exogenous (independent) predictor variables s is denoted as VARMAX(p,q,s). Since some animals move seasonally between the Ngorongoro Crater and the surrounding multiple use areas, the wet and dry season counts do not estimate the same underlying population size. We therefore treat the wet and dry season counts as two separate but possibly correlated variables and use a bivariate VARMAX(p,q,s) model. We allow variation in herbivore numbers in the wet and dry season to depend on the total wet and dry season rainfall in the current year (t) and in the preceding five years (*t*-1,…, *t*-5). The model thus allows the current wet and dry season rainfall components and their lagged values up to five years prior to the current count year to influence the population size of herbivores in the current wet and dry season. The model can also therefore be viewed as a multiple (or distributed) lag regression model. The VARMAX (p,q,s), model we used to forecast the future population dynamics of the five most abundant herbivore species can thus be cast as:

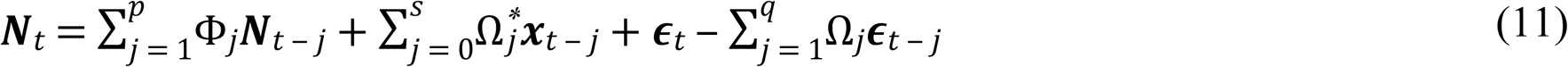

where *N*_*t*_ = (*N*_*wet,t*_, *N*_*dry,t*_)^*T*^are the population sizes of the same species in the wet and dry seasons at time *t, x*_*t*_ = (*wet*_*t* - 0_,…,*wet*_*t* - 5_,*dry*_*t* - 0_,…,*dry*_*t* - 5_)^*T*^ are the wet and dry season rainfall components divided by their long-term means and lagged over 0 to 5 years. *∈*_*t*_ = (*∈*_*wet,t*_,*∈*_*dry,t*_)^*T*^are a two-dimensional vector white noise process. It is assumed that 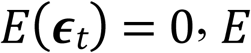 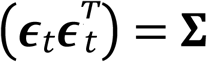 and 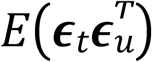 for *t* ≠ *u*. We further assume that *p* and *q* are each equal to either 1 or 2 whereas *s* is set equal to 5. Accordingly, the model can be denoted symbolically as a VARMAX (2,2,5) model. In other words, in order to project the population dynamics of the Ngorongoro large herbivores, we built a model relating the population size of each herbivore species in the current year (t) to the population size in the past one to two years (year t-1 and t-2; i.e., autoregressive process of order *p* = 1 or 2). The model also allows residuals for the current year to depend on the residuals for the previous one to two years (i.e. a moving average process of order *q* =1 or 2). Since herbivore numbers are counted once in the wet season and once in the dry season of each year we did not allow for seasonal variation in the counts.

The VARMAX (p,q,s) model can be represented in various forms, including in state space and dynamic simultaneous equation or dynamic structural equations forms. We used bivariate autoregressive moving average models with the wet and dry season rainfall as the explanatory variables. We tested and allowed for various lags in rainfall so that the models can be characterised as autoregressive and moving-average regression with distributed lags. We also used dead-start models that do not allow for present (current) values of the explanatory variables. We tested for heteroscedasticity in residuals and tested the appropriateness of GARCH-type (generalized autoregressive conditional heteroscedasticity) conditional heteroscedasticity of residuals. We used several information-theoretic model selection criteria as aids to determine the autoregressive (AR) and moving average (MA) orders of the models. The specific criteria we used were the Akaike information criterion (AIC), the corrected AIC (AICc) and the final prediction error (FPE). As additional AR order identification aids, we used partial cross-correlations for the response variable, Yule-Walker estimates, partial autoregressive coefficients and partial canonical correlations. Parameters of the selected full models were estimated using the maximum likelihood (ML) method. Roots of the characteristic functions for both the AR and MA parts (eigenvalues) were evaluated for the proximity of the roots to the unit circle to infer evidence for stationarity of the AR process and inevitability of MA process in the response series.

The adequacy of the selected models was assessed using various diagnostic tools. The specific diagnostic tools we used are the following. 1) Durbin-Watson (DW) test for first-order autocorrelation in the residuals. 2) Jarque-Bera normality test for determining whether the model residuals represent a white noise process by testing the null hypothesis that the residuals are normally distributed. 3) F tests for autoregressive conditional heteroscedastic (ARCH) disturbances in the residuals. This F statistic tests the null hypothesis that the residuals have equal covariances. 4) F tests for AR disturbance computed from the residuals of the univariate AR(1), AR(1,2), AR(1,2,3) and AR(1,2,3,4) models to test the null hypothesis that the residuals are uncorrelated. 5) Portmanteau test for cross correlations of residuals at various lags. Final forecasts and their 95% confidence intervals were then produced for the animal population size series for each of the five most common species in each season for lead times running from 2013 up to 2100.

In the table of the parameter estimates for the bivariate VARMAX (2,2,5) model fitted to the two time series of herbivore population size in the wet and dry seasons (Table S1), the five lagged dry and wet season rainfall components (rightmost column labelled variable) for the current year (year *t*) up to five years prior to the current year (years *t*-1,…, *t*-5) are denoted by dry (*t*),…, dry (*t*-5) and wet (*t*),…, wet (*t*-5), respectively. Analogously, for the dry season counts, the autoregressive process of order 2 is denoted by, e.g., wildebeest_dry_(*t*-1) and wildebeest_dry (*t*-2) while the moving average process of order 2 by e1(*t*-1) and e2 (*t*-2). A parallel notation is used for the wet season counts. The estimated regression coefficients (estimate) for the parameters associated with each of these variables plus the intercept (Const1), the standard errors of the estimates and a *t*-test (t-value) of the null hypothesis that each coefficient is not significantly different from zero (Pr >|*t*|) are also provided in Table S1. Furthermore, the estimated roots of the autoregressive (Table S2) and moving average (Table S3) processes are provided. It is important to note that the population of each herbivore species in the wet season of the current year depends not only on its lagged values in the preceding one to two years and on the current and past values of rainfall but also on the population of the same herbivore species in the dry season lagged over the past one to two years. The same applies to the population of each herbivore species in the current dry season. This interdependence of the two series on each other is made possible because of the bivariate nature of the VARMAX (p,q,s) model. This model was fitted to the population counts of the herbivores for the wet and dry seasons for the period 1964-2012 based on historic station rainfall data for 1963 to 2012. Note that the historic total wet season rainfall component was divided by its mean for use in the model. The same was done for the total dry season rainfall component. Future forecasts were then produced by supplying the projected wet and dry season rainfall values, each divided by its mean, for Ngorongoro for 2013 to 2100.

Several univariate model diagnostics were used to extensively assess how well the selected bivariate VARMAX (p,q,s) model fitted the count data (Tables S4-S7). The first model diagnostic tool, the Portmanteau Test for Cross Correlations of Residuals (Table S4) was significant, considering only up to lag 5 residuals. This test of whether the residuals are white noise residuals (i.e. uncorrelated) based on the cross correlations of the residuals, suggests that the residuals were apparently correlated, when only up to lag 5 residuals are considered. Even so, results of the univariate model ANOVA diagnostics suggest that the models for both the dry and wet season counts were highly significant and had high predictive power (*r*^2^, Table S5). Results of the Univariate Model White Noise Diagnostics (Table S6) suggest that the residuals are normally distributed (Jarque-Bera normality test) and have equal covariances (ARCH (1) disturbances test). The Univariate AR Model Diagnostics indicate that the residuals are uncorrelated, contrary to the finding of the multivariate Portmanteau test (Table S7). The modulus of the roots (eigenvalues) of the AR characteristic polynomial are less than 1 suggesting that the series are stationary. These tests suggests that the fitted models are reasonable. The log-transformed animal count totals, rainfall deviates, projected rainfall and forecast animal count totals (log scale) are provided in S4 Data. The SAS program codes used to analyze the rainfall data are provided in S1 Text while the code for analyzing the animal counts is provided in S2 Text.

## Results

### Rainfall

Rainfall can be subdivided into the dry and wet season components. The dry season occurs from June to October whereas the wet season occurs from November to May. The wet season rainfall component is strongly bimodal, with the two modes corresponding to peaks in the long rains and the short rains. The major peak in rainfall occurs in April during the long rains (January-May) whereas the minor peak occurs in December during the short rains (November-December, Fig 2a). The total monthly rainfall averaged 78.3 ± 84.2 mm and was highly variable (%CV = 107.5%) during 1963-2014 (Fig 2a). The total annual rainfall averaged 937.5 ± 300.7 mm during 1963-2014 (Fig. 2b) out of which the wet season rainfall (851.7 ± 297.3 mm) contributed 90.9% (Fig 2c) and the dry season rainfall (85.5 ± 65.2 mm) a mere 10.1% (Fig 2d). There were also considerable interannual variations in the annual, wet and dry season rainfall components (Figs 2b-d). Smoothing of the time series of the total monthly rainfall exposed substantial variation with periods of below-average rainfall centered around 1966, 1975, 1980 and 1999 (Fig S1).

**Fig 2.**
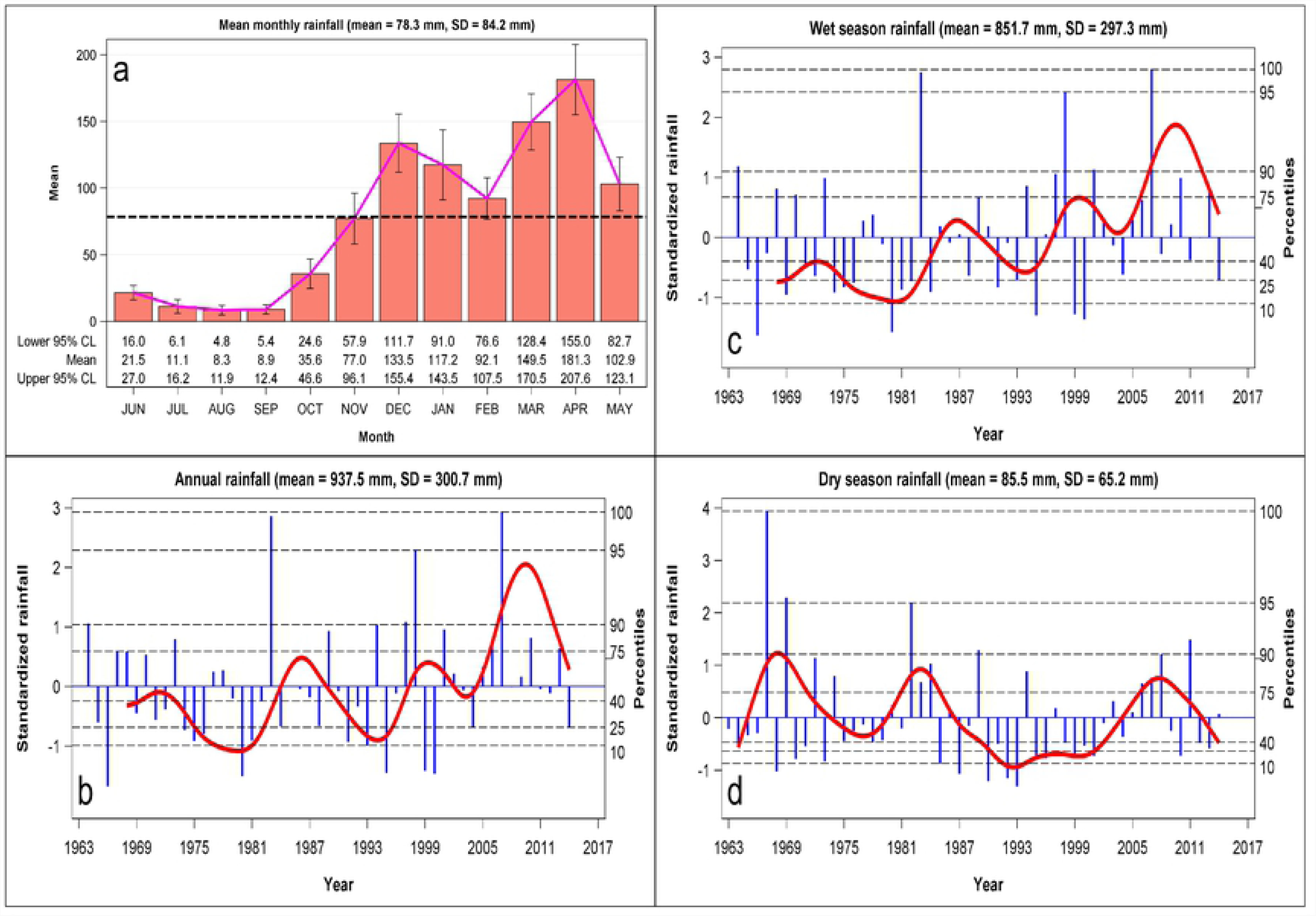
The distribution of a) total monthly rainfall (mean ± 1sd = 78.3 ± 84.2 mm) across months in the Ngorongoro Crater National Park averaged over 1963-2014 and the interannaul variation in standardized deviations of the b) annual rainfall (mean ± 1SD =937.5 ± 300.7 mm), c) wet season rainfall (mean± 1SD =851.7 ± 297.3 mm), and d) dry season rainfall (mean± 1SD =85.5 ± 65.2 mm) in the Ngorongoro Crater during 1963-2014. The vertical needles are the standardized deviates, the solid curves are the 5-year (annual and wet season) and 2-year (dry season) moving averages and the dashed horizontal lines are percentiles of the frequency distributions of the rainfall deviates.

Analysis of the annual rainfall showed that extreme droughts occurred in 1966, 1980, 1993, 1995, 1999 and 2000 while severe droughts were recorded in 1974-1976, 1981, 1991, 2004 and 2014. Further, the extremely wet years were 1983 and 2007 whereas very wet years were 1964, 1997 and 1998. Analysis of the wet season rainfall identified the same extreme and severe droughts and very wet years as the annual rainfall did (Table S8, Fig S2). In addition, the wet season of 1969 experienced an extreme drought while the 1982 wet season was a severe drought. The dry seasons of 1968, 1985, 1987, 1990, 1992 and 1993 were extremely dry and the dry seasons of 1970, 1973, 1995, 1996, 1999, 2001 and 2010 were severe droughts. By contrast the dry seasons of 1967, 1969, 1982, 1989 and 2011 were either extremely wet or very wet (Table S8, Fig S2).

There were significant quasi-cyclic oscillations in the three rainfall components with approximate cycle periods of 4.64, 4.64 and 2.47 years for the annual, wet season and dry season rainfall components, respectively, based on spectral analysis (Table S9, Fig S3). Based on the unobserved components model (UCM), the oscillations in the annual, wet season and dry season rainfall components had dominant cycle periods of 4.83, 3.82 and 2.45 years, respectively (Table S10, Fig S4). In addition, there were secondary cycles in the wet and dry season rainfall components with approximate cycle periods of 2.2 years for the wet season component and 11.3 years for the dry season component (Table S10, Figs S4). The estimated damping factors for the cycles were all less than 1 except for the cycle for the annual rainfall component with a period of 4.83 years and the cycle with a period of 2.2 years for the wet season rainfall component both of which had damping factors equal to 1 (Table S10, Fig S4). The two cycles with damping factors equal to 1 are persistent whilst the remaining cycles with damping factors smaller than 1 are transient.

The disturbance variances for the irregular components for the wet and dry season rainfall, but not for the annual rainfall, were close to zero and statistically insignificant. This implies that the irregular components for the two seasonal rainfall components were deterministic whereas the irregular component for the annual rainfall was stochastic. Moreover, the estimated disturbance (error) variances for the cyclical components were significant for the 3.83-year cycle for the wet season and for both cycles for the dry season but not for the 4.83-year cycle for the annual rainfall (Table S10, Figs S4). These features jointly imply that the 4.83-year cycle identified for the annual rainfall is persistent and deterministic whereas the cycles identified for both the wet and dry season rainfall are stochastic and transient (Table S10, Figs S4). Even so, significance analysis of the disturbance (error) variances of the cyclical components in the model at the end of the estimation span indicate that the disturbance variances for the cycle in the annual rainfall component and both cycles in the wet season rainfall component were significant but those for the two cycles in the dry season rainfall component were insignificant (Table S11). Since the 4.83-cycle in the annual rainfall component is deterministic the additional significant test result means that the annual cycle is indeed significant. The significant disturbance variances for the two stochastic cycles in the wet season rainfall component (Table S11) applies only to the part of the time series of wet season rainfall near the end of the estimation span.

The disturbance terms for the level component for all the three rainfall components were significant only for the wet season but not for the annual or dry season rainfall. As well, the slope component was significant only for the wet season rainfall (Table S11, Fig S4). This implies that, of the three rainfall components, only the wet season rainfall increased systematically over time in Ngorongoro (Table S11, Fig S4). The smoothed rainfall cycles in the three rainfall components further reinforce the conclusion that the oscillation in annual rainfall is persistent and deterministic whereas the oscillations in the wet and dry season rainfall are transient and stochastic (Fig S4).

### Projected rainfall and temperatures

The projected annual rainfall showed no evident systematic trend under all the three scenarios. However, the general average rainfall level is consistently and substantially higher under the RCP2.6 than the RCP4.5 and 8.5 scenarios. The RCP4.5 and 8.5 scenarios have comparable average levels but RCP4.5 is expected to receive somewhat more rainfall. Notably, rainfall shows marked inter-annual variation characterized by sustained quasi-cyclic oscillations during 2006-2100 regardless of scenario (Fig S5).

The minimum and maximum temperatures are expected to rise during 2006-2100, on average, by 1, 2 and 6 °C under the RCP2.6, 4.5 and 8.5 scenarios, respectively. Consequently, the average maximum temperature is expected to increase during 2006-2100 from 23 to 24 °C under RCP2.6, 24 to 26 °C under RCP4.5 and 23 to 29 °C under RCP8.5. The average minimum temperature is similarly anticipated to rise during 2006-2100 from 14 to 15 °C under RCP2.6, 14 to 16 °C under RCP4.5 and 14 to 20 °C under RCP8.5 (Fig S5).

### Changes in vegetation composition and structure

From 1966/67 [14] to 1995 [38] there have been significant changes in the structure of the major and secondary herbaceous species. In 1966/67 the Crater floor was dominated by short grass herbaceous species. By 1995, most of the short grasslands had been replaced by mid to tall plant species.

### Historic herbivore population dynamics

The population size of wildebeest, zebra, Thomson’s gazelle, Grant’s gazelle, kongoni (Coke’s hartebeest), and black rhino increased from 1964 to a peak around 1974-1976 and then declined thereafter in both the wet and dry seasons. Eland and waterbuck had a general downward trend from the early 1970’s. Zebra numbers increased again from 1995 to 2012 whereas Grant’s gazelle and kongoni numbers in the dry season increased again from 1995 to 2000 before declining further (Fig 3). In stark contrast to the other species, numbers of buffalo increased markedly following the removal of Maasai livestock from the Crater in 1974. Elephant and ostrich numbers have similarly increased in the Crater, with substantial increase apparent in ostrich numbers following the extreme 1993 dry season drought (Fig 3). Buffalo, eland, elephant and black rhino were more abundant in the Crater in the wet than the dry season. There were far more eland and black rhino in the Crater in the wet season compared to the dry season in the 1970s than in the 2000s. Conversely, there were far more buffalo and elephants in the Crater in the wet season compared to the dry season in the 2000s than there were in the 1970s (Fig 3). Zebra were the only species to have maintained similar population sizes from 1964 to 2012.

**Fig 3.**
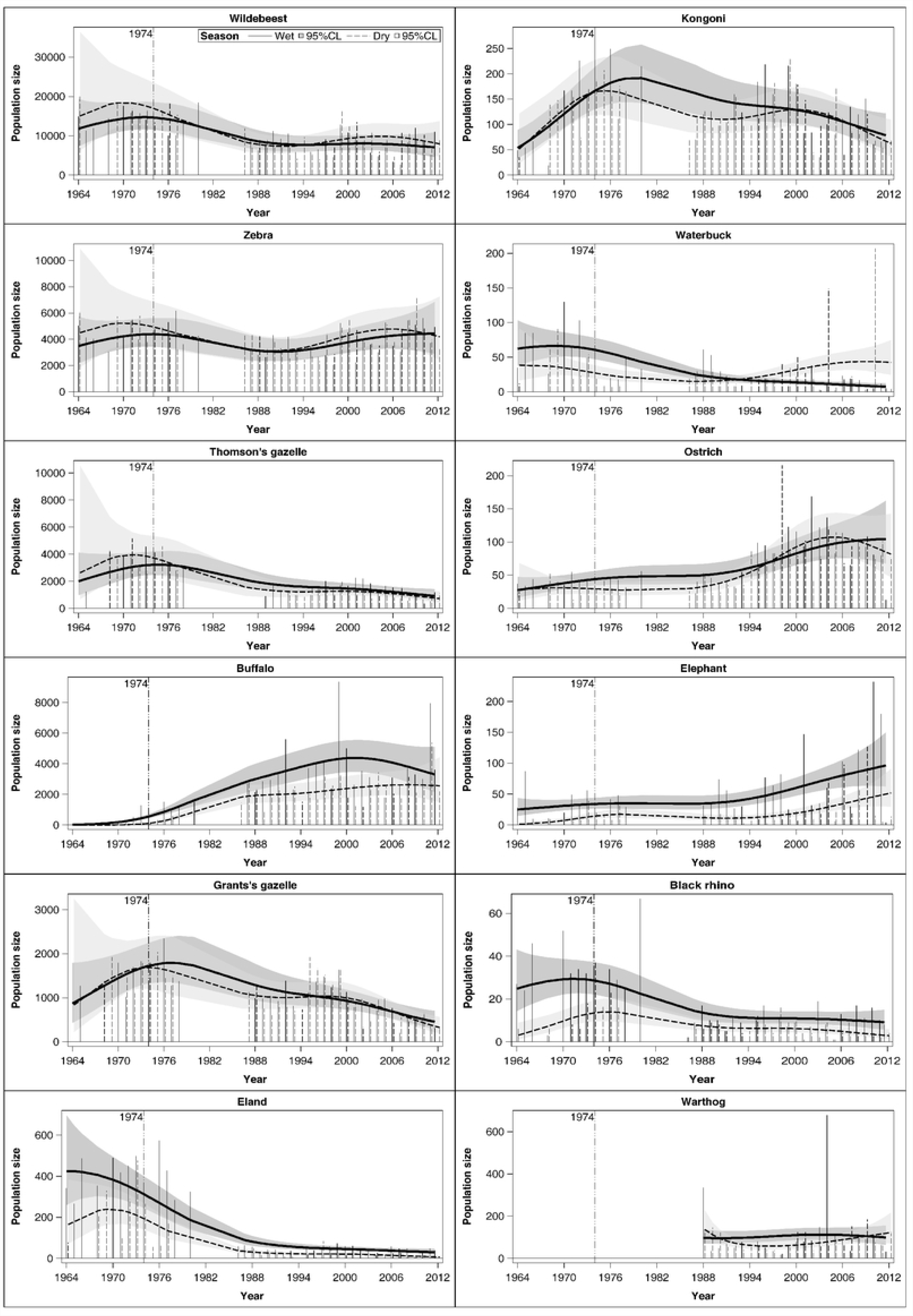
Trends in the population sizes of the 12 most common large herbivore species in the Ngorongoro Crater in the wet and dry seasons from 1964 to 2012. The vertical needles denote wet season (solid) and dry season (dashed) count totals. Thick solid and dashed curves denote the fitted wet season and dry season trend curves. The shaded regions are the 95% point wise confidence bands.

Comparisons of the expected population sizes between 1964 and 1974 as well as between 1974 and 2012 based on constructed spline effects showed that while some species increased significantly over time, others did not, or even declined. Species that increased but not significantly between 1964 and 1974 in the wet season were wildebeest, Grant’s gazelle, waterbuck and ostrich (Table S12, Fig 3). Only buffalo, Thomson’s gazelle and kongoni numbers increased significantly between 1964 and 1974 in the wet season. Species that decreased in numbers but not significantly between 1964 and 1974 in the wet season were zebra, eland, elephant and black rhino. Between 1974 and 2012, the numbers of Thomson’s gazelle, Grant’s gazelle, black rhino, eland, kongoni and waterbuck decreased significantly in the wet season. In the same season and period, the numbers of buffalo and elephant increased significantly. Zebra, wildebeest, and ostrich had no significant change (Table S12, Fig 3). In the dry season, by contrast, numbers of some species either increased significantly between 1964 and 1974 (buffalo, elephant, eland, kongoni), increased but not significantly (waterbuck) or decreased but not significantly (wildebeest, zebra, Thomson’s gazelle, Grant’s gazelle, ostrich). However, between 1974 and 2012 in the dry season, numbers of some species either increased significantly (buffalo, ostrich), increased but not significantly (waterbuck), decreased significantly (Thomson’s gazelle, Grant’s gazelle, black rhino, eland, kongoni), or decreased but not significantly (wildebeest, zebra, elephant, Table S12, Fig 3).

### Herbivore biomass dynamics

Herbivore biomass in the wet season was initially dominated by wildebeest, followed by zebra. Following the eviction of the Maasai and their livestock from the Crater in 1974, buffalo biomass increased relative to wildebeest and zebra to a peak during 1999-2000. After the 1999-2000 drought, the biomass of buffalo and the other herbivore species declined to the pre-drought levels. Nevertheless, wildebeest still makes a smaller contribution to the total biomass currently than they did before cattle left the Crater and buffalo numbers were still low (Fig 4a). The relative increase of buffalo biomass compared to wildebeest and zebra was also apparent in the dry season biomass (Fig 4b).

**Fig 4.**
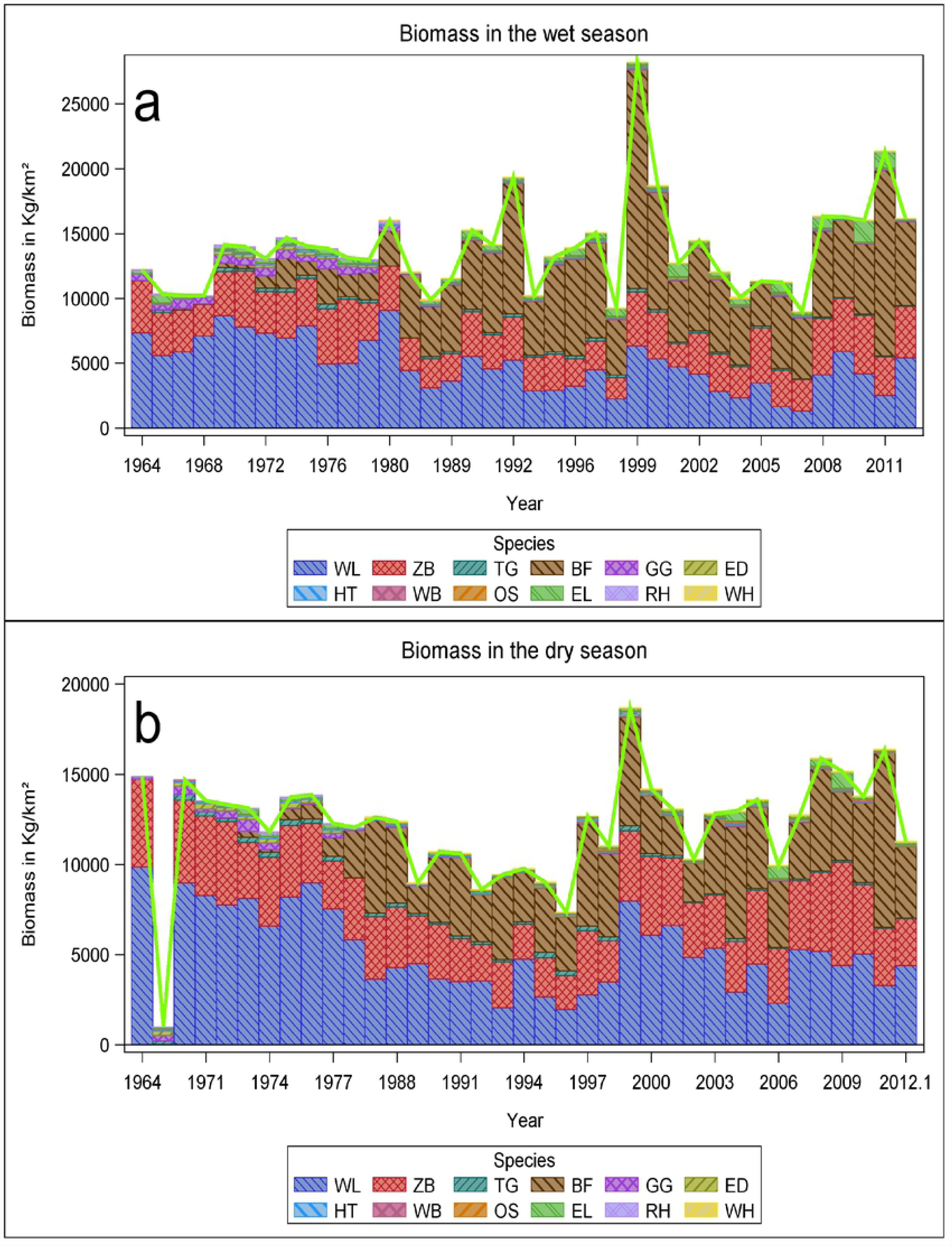
Temporal trends in the cumulative total biomass (kg) of the 12 most common large herbivore species in the Ngorongoro Crater during a) the wet season and b) the dry season during 1964 to 2012. The unit weights (kg) are 1725, 816, 450, 340, 200, 160, 125, 123, 114, 45, 40 and 15 for elephant, rhino, buffalo, eland, zebra, waterbuck, kongoni, wildebeest, ostrich, warthog, Grant’s gazelle and Thomson’s gazelle, respectively. Note that wildebeest and zebra were not counted in the dry season of 1968. In years when multiple surveys were done in the same season (e.g., the wet season of 1966 or 1970), only the survey with the maximum count was used to calculate biomass.

The total herbivore biomass trends in the Crater have been dynamic and relatively stable. During the dry season from 1964 to 1974 there was no significant change and this trend was also non-significant for the dry season from 1974 to 2011 (Table 2). This scenario of a non-significant trend from 1964 to 1974 and again from 1974 to 2011 was also consistent for the wet season (Table 2).

**Table 1.** Changes in vegetation structure from 1966/67 to 1995.

| Major Herbaceous Species (ha) |  |  |
| --- | --- | --- |
|  | Herlocker | Chuwa |
| Short | 20868 | 10626 |
| Mid | 2110 | 9540 |
| Mid-tall | 5794 | 4506 |
| Tall | 1377 | 4227 |
| Secondary Herbaceous Species (ha) |  |  |
| Short | 18584 | 4794 |
| Mid | 2812 | 3905 |
| Mid-tall | 2004 | 9096 |
| Tall | 1317 | 2258 |

**Table 2.** The expected aggregate biomass in the wet and dry seasons of 1964, 1974 and 2011 and the difference between the 1964 vs 1974 and 1974 vs 2011 estimates and test of significance of their difference based on constructed penalized cubic B-splines.

| Statement Number | Label | Estimate | Standard Error | DF | t Value | Pr > \|t\| | Adjustment | Adj P |
| --- | --- | --- | --- | --- | --- | --- | --- | --- |
| 1 | Dry Season at time=1964 | -63.3804 | 265.19 | 2.326 | -0.24 | 0.8306 |  | . |
| 2 | Dry Season at time=1974 | -41.5788 | 146.46 | 2.349 | -0.28 | 0.7996 |  | . |
| 3 | Dry Season at time=2011 | 8.0933 | 0.1129 | 47.44 | 71.67 | <0.0001 |  | . |
| 5 | Wet Season at time=1964 | -64.2763 | 265.14 | 2.326 | -0.24 | 0.8282 |  | . |
| 6 | Wet Season at time=1974 | -41.5482 | 146.46 | 2.349 | -0.28 | 0.7998 |  | . |
| 7 | Wet Season at time=2011 | 8.4434 | 0.1336 | 50.95 | 63.2 | <0.0001 |  | . |
| 4 | Diff for Dry Season at time= 1964 vs time= 1974 | 21.8017 | 123.24 | 2.309 | 0.18 | 0.8739 | Simulated | 0.9479 |
| 4 | Diff for Dry Season at time= 1974 vs time= 2011 | 49.6721 | 146.46 | 2.349 | 0.34 | 0.7625 | Simulated | 0.8474 |
| 8 | Diff for Wet Season at time= 1964 vs time= 1974 | 22.7281 | 123.16 | 2.309 | 0.18 | 0.8686 | Simulated | 0.9447 |
| 8 | Diff for Wet Season at time= 1974 vs time= 2011 | 49.9916 | 146.46 | 2.348 | 0.34 | 0.761 | Simulated | 0.8463 |

### Relationship between herbivore population size and rainfall

Herbivore population size was correlated with rainfall in both the wet and dry seasons. The particular rainfall component most strongly correlated with population size as well the specific functional form of the relationship both varied with species and season (Figs 5 and 6, Tables S13-S14). In the wet season, population size was most tightly correlated with 1) 6-year moving averages of the wet season rainfall (wildebeest, zebra, buffalo, eland, kongoni, waterbuck, ostrich, elephant, black rhino), 2) 6-year moving average of the annual rainfall (Thomson’s and Grant’s gazelles), or 3) the current annual rainfall (warthog). In the dry season, population size had the strongest correlation with 1) 6-year moving average of the wet season rainfall (Thomson’s and Grant’s gazelle, buffalo, waterbuck, ostrich), 2) 5-6-year moving average of the dry season rainfall (wildebeest, zebra, warthog), 3) 6-year moving average of the annual rainfall (eland, kongoni), or 4) 3-4-year moving average of dry season rainfall (elephant, black rhino, Figs 5 and 6, Tables S13-S14). The dependence of population size on rainfall followed three general patterns. The first pattern is characterized by a decline in population size with increasing rainfall and is shown by wildebeest, eland, kongoni, waterbuck and black rhino in the wet season, and Thomson’s gazelle, Grant’s gazelle and waterbuck in the dry season. The second pattern consists of an increase in population size with increasing rainfall and is shown by zebra, buffalo, ostrich and elephant in the wet season and wildebeest, zebra, buffalo, ostrich and warthog in the dry season. The third and last pattern is characterized by a humped relationship between population size and rainfall in which population size peaks at intermediate levels of rainfall and is shown by Thomson’s and Grant’s gazelles and warthog in the wet season and eland, kongoni, elephant and black rhino in the dry season (Figs 5 and 6, Tables S13-S14).

**Fig 5.**
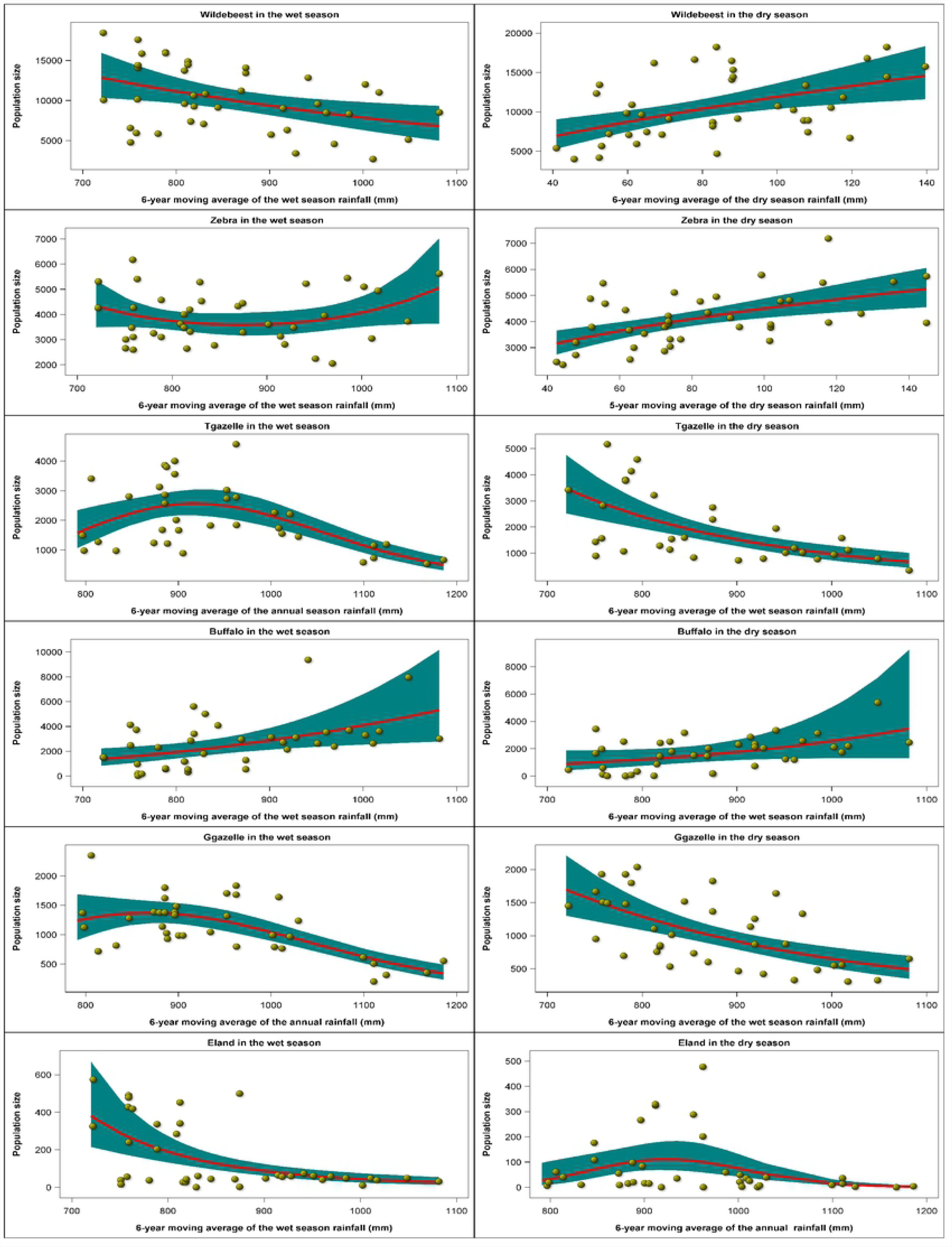
The selected best regression relationships between the wet season and dry season count totals of wildebeest, zebra, Thomson’s gazelle, buffalo, Grant’s gazelle, and eland and the moving averages of the annual, wet season and dry season rainfall components for the Ngorongoro Crater during 1964-2012.

**Fig 6.**
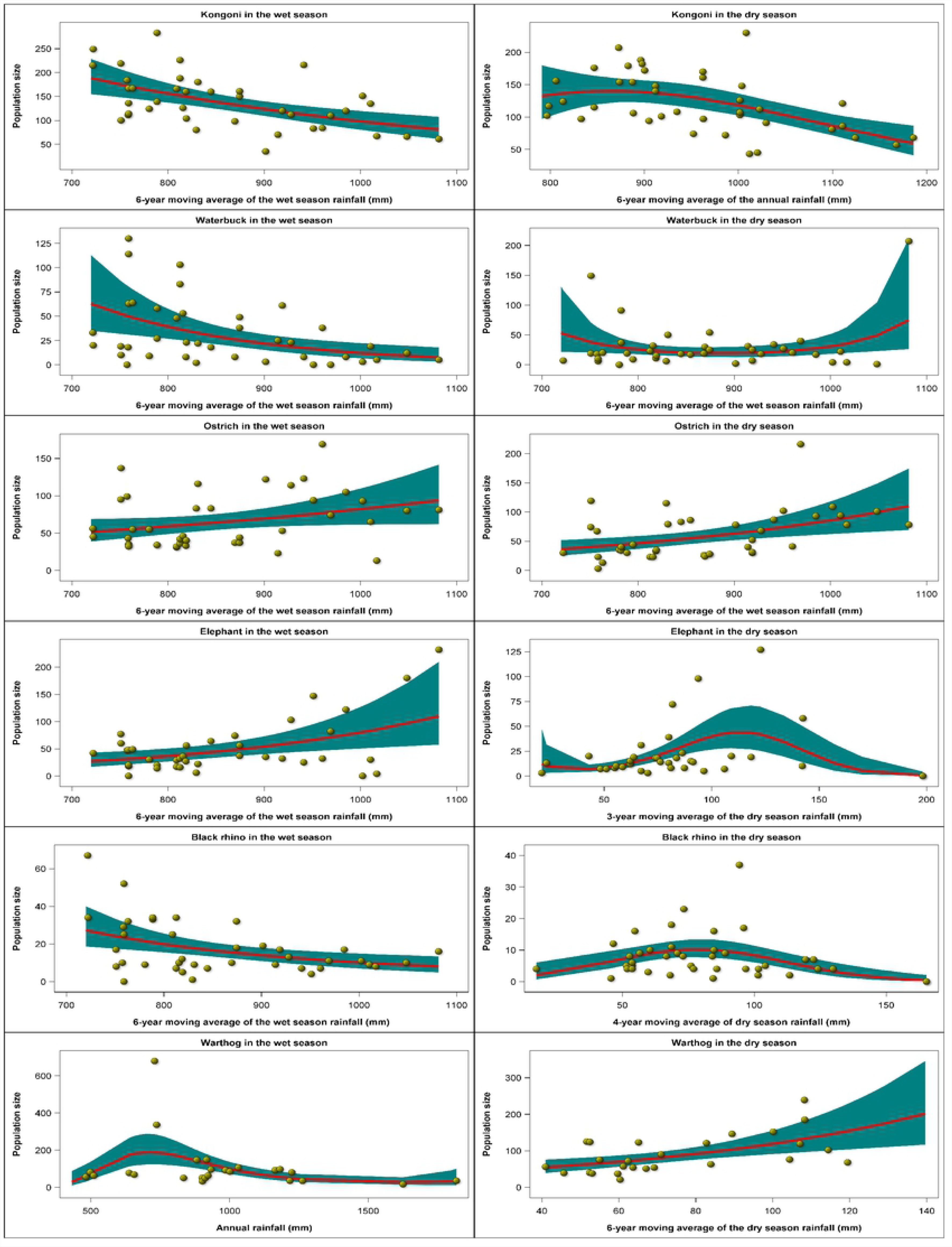
The selected best regression relationships between the wet season and dry season count totals of kongoni, waterbuck, ostrich, elephant, black rhino and warthog and the moving averages of the annual, wet season and dry season rainfall components for the Ngorongoro Crater during 1964-2012.

### Projected herbivore population dynamics

The projected ungulate population dynamics should mirror the pronounced and sustained oscillations in the projected rainfall. Further, large-sized herbivores dependent on bulk, low-quality forage should prosper under the wet and cooler conditions expected under RCP2.6. Likewise, small-sized herbivores requiring high-quality forage should thrive under the relatively low rainfall and warmer conditions anticipated under RCP4.5 and 8.5. The warmer temperatures expected under RCP8.5 than under RCP4.5 imply that conditions should be most arid under this scenario.

The projected population trajectories suggest that under the RCP2.6 scenario, buffalo numbers will likely continue to increase after 2012, albeit at a decelerating rate, towards 7000-11000 animals by 2100 (Fig 7). But the Crater buffalo population is likely approaching its upper bound of about 4000 animals and will likely fluctuate about this number (4000) till 2100 under the RCP4.5 and 8.5 scenarios regardless of season (Fig 7). As expected, the population of this large-sized bulk grazer is projected to be highest on average under RCP2.6, least under RCP8.5 and intermediate under RCP4.5 for both the wet and dry seasons (Fig 7).

**Fig 7.**
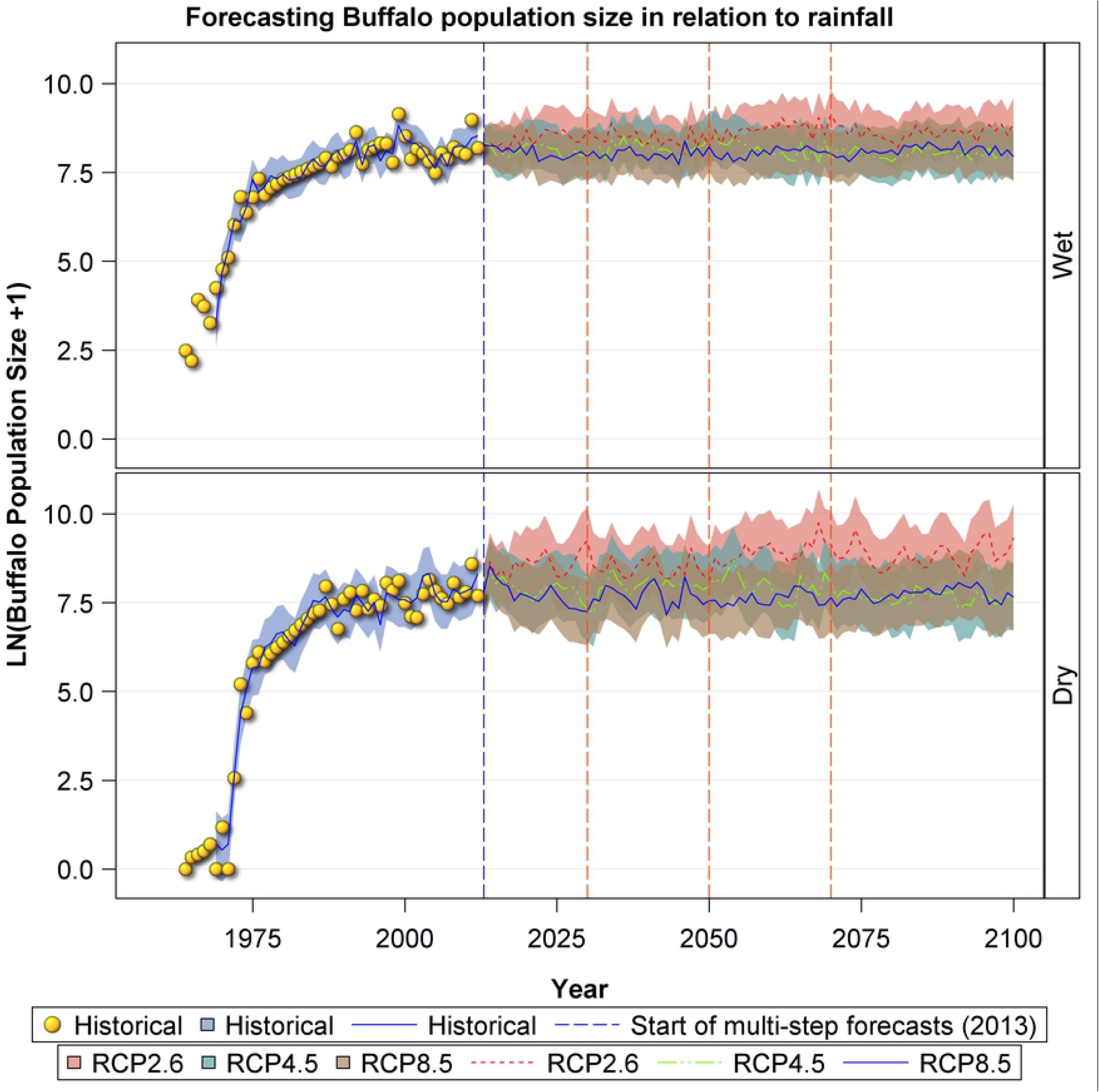
Historic and projected population size of buffalo in the Ngorongoro Crater during the wet and dry seasons based on the three climate change scenarios RCP2.6, RPC4.5 and RCP8.5.

For wildebeest, the projected trajectories suggest strong and sustained oscillations in population size under all the three scenarios and both seasons, reflecting the strong projected rainfall oscillations (Fig 8). The oscillatory population dynamics in both the wet and dry seasons exhibited by wildebeest reveal extended periods of population increase followed by prolonged periods of persistent population declines. Nevertheless, there are also discernible differences in the projected population trajectories under the three climate change scenarios. The projected wildebeest population trajectories suggest that the population will continue to fluctuate widely between 5000 and 15000 animals in all the scenarios and seasons. It is only under the RCP2.6 scenario that the dry season population shoots beyond 20000 animals around 2070 and 2090 (Fig 8). In the wet season, the projected average wildebeest abundance is highest under RCP4.5, intermediate under RCP8.5 and lowest under RCP2.6. In the dry season, however, wildebeest abundance is highest on average under RCP2.6, intermediate under RCP4.5 and lowest under RCP8.5 (Fig 8).

**Fig 8.**
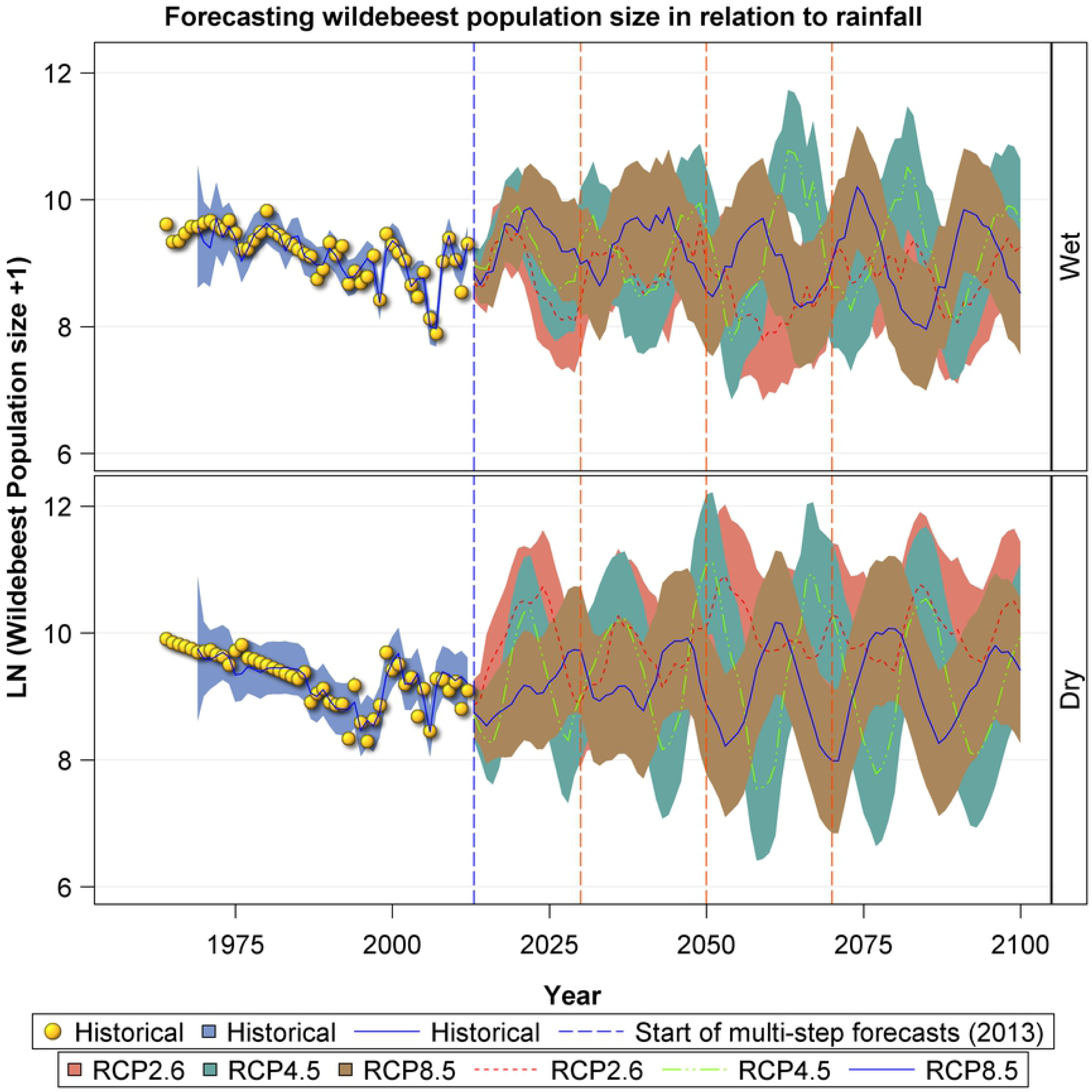
Historic and projected population size of wildebeest in the Ngorongoro Crater during the wet and dry seasons based on the three climate change scenarios RCP2.6, RCP4.5 and RCP8.5.

The zebra population trajectories also reveal striking oscillations in population size under all the three scenarios, a general increase in population size under RCP2.6 scenario in both seasons and a decrease and then increase in the RCP8.5 scenario in the wet season (Fig 9). The zebra population size is projected to decline in the long term under the RCP4.5 scenario in both seasons and the RCP8.5 scenario in the dry season (Fig 9). In general, zebra will perform the best under RCP2.6 and the worst under RCP8.5. The performance of zebra under RCP4.5 will be intermediate between RCP2.6 and 8.5 from 2006 to around 2070 after which it will drop below that expected under RCP8.5 (Fig 9).

**Fig 9.**
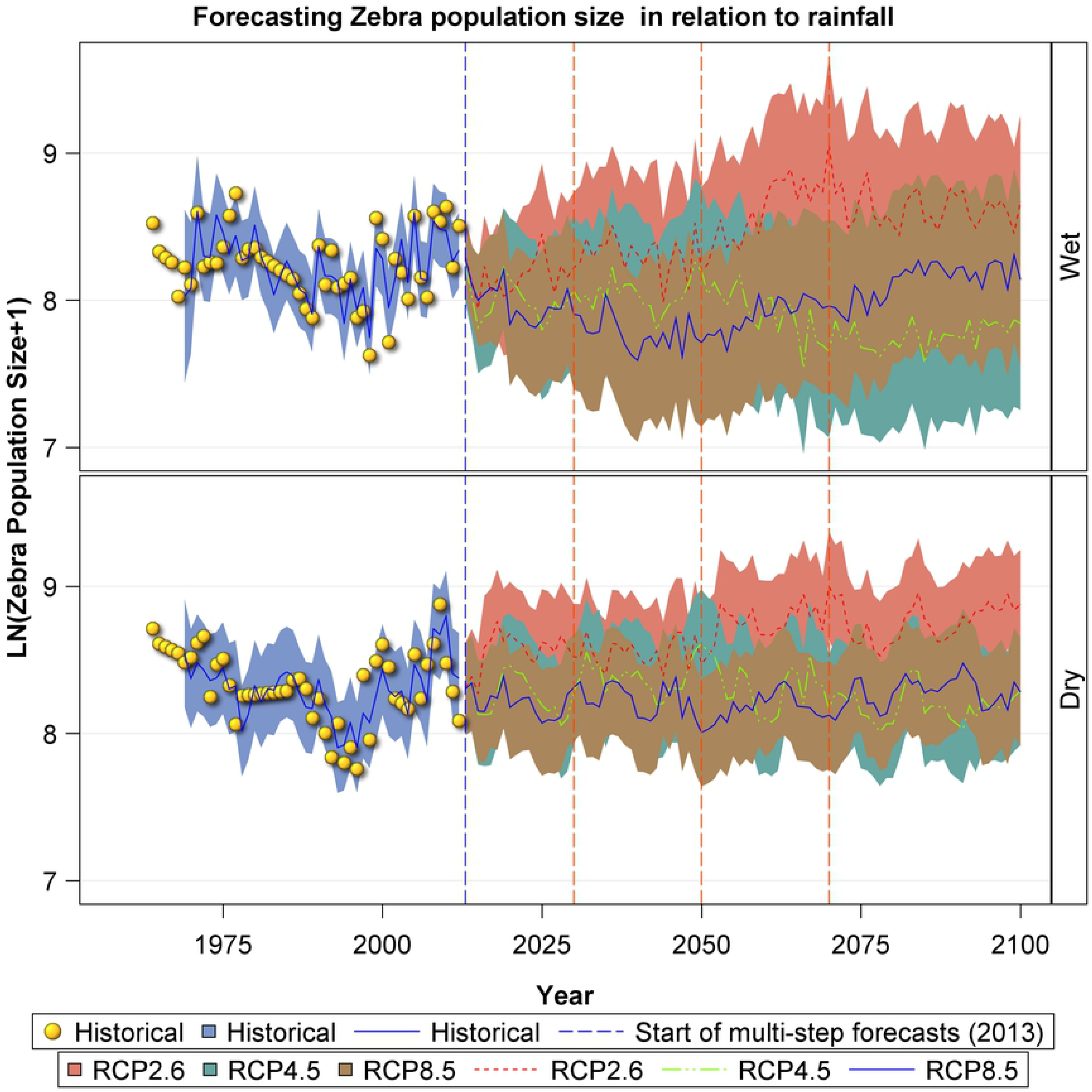
Historic and projected population size of zebra in the Ngorongoro Crater during the wet and dry seasons based on the three climate change scenarios RCP2.6, RCP4.5 and RCP8.5.

The decline observed in historic Thomson’s gazelle numbers is projected to be persistent and to remain below the peak attained historically around 1974 under all scenarios and both seasons (Fig 10). Besides the general decline, Thomson’s gazelle numbers are projected to show persistent and marked oscillations irrespective of scenario or season. As predicted by their small body size and selective grazing, Thomson gazelles will likely perform the best under RCP8.5 with the least rainfall, intermediately under RCP4.5, and the worst under RCP2.6 (Fig 10).

**Fig 10.**
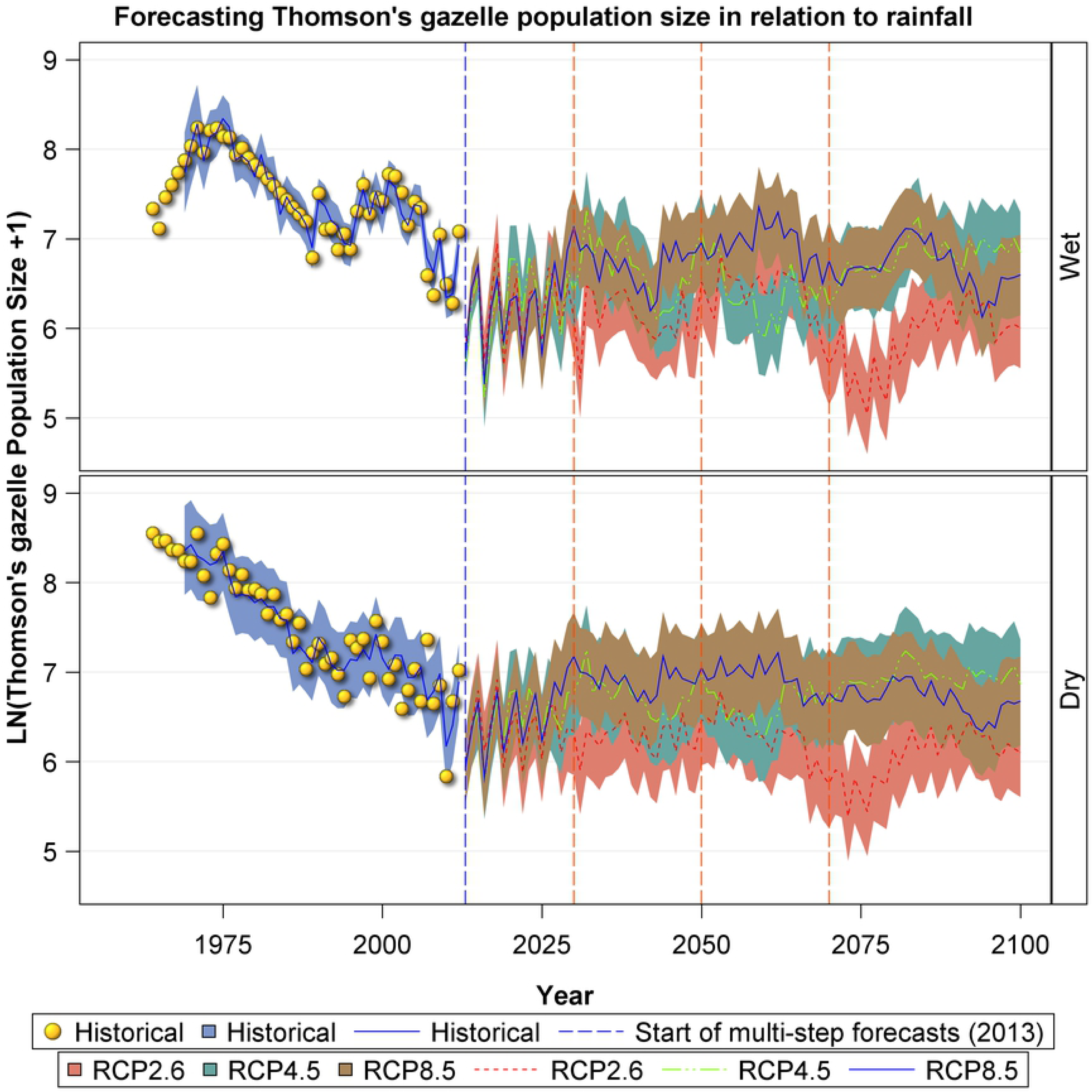
Historic and projected population size of Thomson’s gazelle in the Ngorongoro Crater during the wet and dry seasons based on the three climate change scenarios RCP2.6, RCP4.5 and RCP8.5.

As with Thomson’s gazelles, the projected population trajectories for Grant’s gazelle show marked and sustained oscillations (Fig 11). Despite these persistent oscillations, Grant’s gazelle numbers will likely remain lower than the historically attained peak numbers around 1974-1976. Moreover, the declining trend in Grant’s gazelle numbers is projected to be replaced by an increasing trend after some time under the RCP4.5 and 8.5 scenarios for both seasons. Even, so Grant’s gazelle numbers, are less likely to increase up to the highest historically recorded numbers around 1974-1976 (Fig 11). Consistent with their small body size and selective grazing, Grant’s gazelles will also likely flourish the best under RCP8.5 with the least rainfall, intermediately under RCP4.5, and the worst under RCP2.6 (Fig 11).

**Fig 11.**
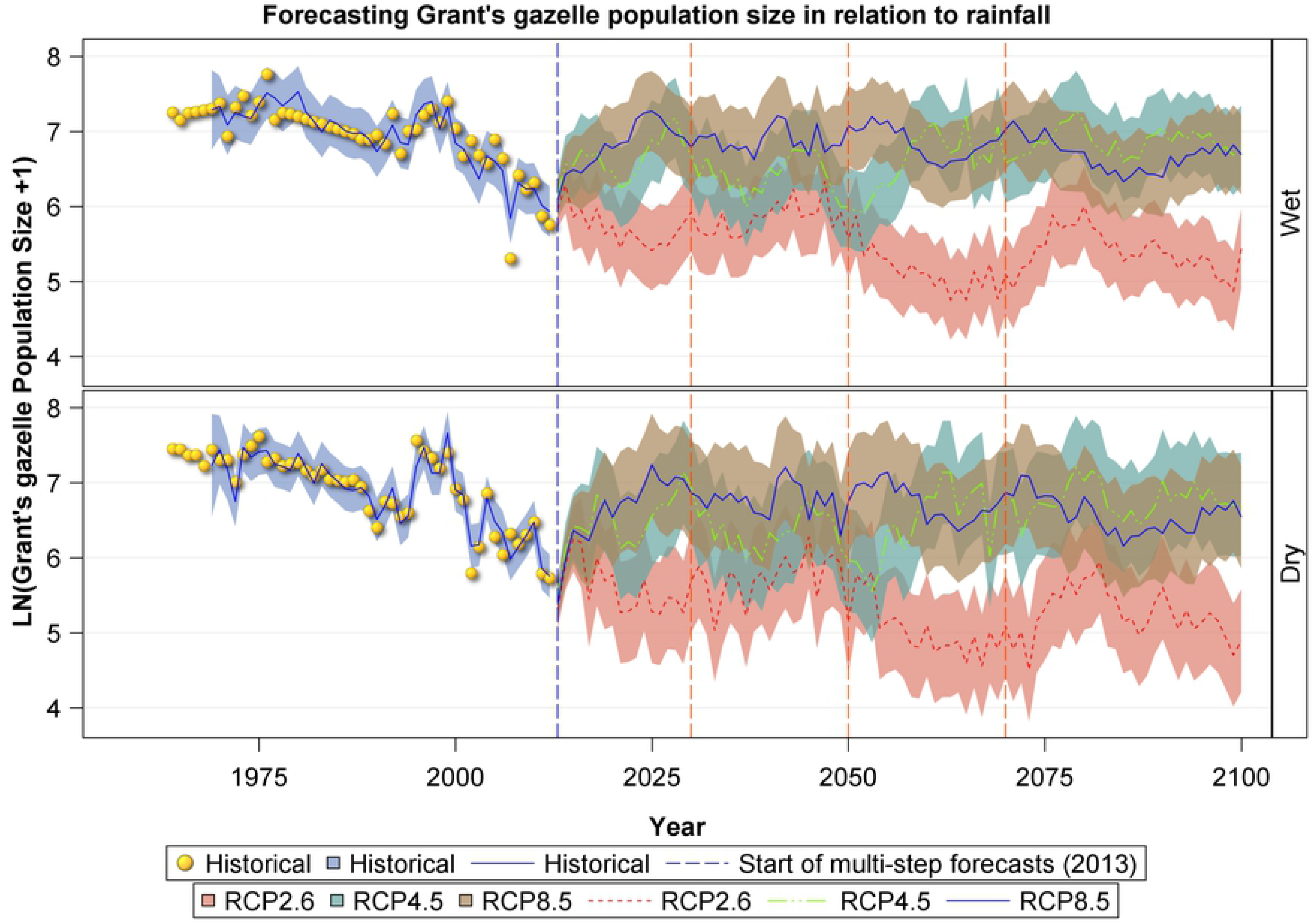
Historic and projected population size of Grant’s gazelle in the Ngorongoro Crater during the wet and dry seasons based on the three climate change scenarios RCP2.6, RCP4.5 and RCP8.5.

## Discussion

### Rainfall

Drought is a recurrent feature of the Ngorongoro Conservation Area. The annual rainfall shows evident persistent and deterministic quasi-periodic oscillation with a cycle period of about 5 years. Oscillations in the wet and dry season rainfall were stochastic and transient. The quasi-cyclic oscillations in annual, wet and dry season rainfall were statistically significant. The oscillations are associated with recurrent severe droughts that cause food scarcity and hence nutritional stress for the large herbivores. The wet season rainfall increased systematically in Ngorongoro between 1964 and 2014 but the annual or dry season rainfall did not increase. The oscillations in rainfall imply that the large herbivores are exposed to above average food supply for about 2.5 years and to below average food supply for the subsequent 2.5 years. The rainfall patterns also imply that portions of the Crater may be waterlogged or flooded during the high rainfall years. High rainfall supports above-average production of plant biomass. But the forage produced during high rainfall years is likely to be of low quality due to the dilution of plant nutrients. Predation risk for herbivores is also likely to rise due to poor visibility associated with tall grass growth during periods of high rainfall [60].

### Long-term Vegetation Trends

Vegetation maps from before and after the removal of pastoralists and their livestock indicate that major changes in vegetation structure occurred. Maasai pastoralists manage their grazing areas with movement of livestock and fire [16, 61,62]. This type of range management selects for shorter grasses and more palatable species [63,64]. The 1995 vegetation map shows that there was a significant change in the vegetation structure of the Crater floor, such that there was a decrease in the availability of short grasses and an increase in medium and tall grassland.

### Historic herbivore population dynamics

Temporal variation in herbivore numbers in the Crater followed four general patterns. First, buffalo, elephant and ostrich numbers increased significantly in the Crater from 1974-2012. The transition of the Crater grasslands to a majority of the area being mid to tall-grass would have favored Cape buffalo reproduction and survivorship. The increase in ostrich and elephant numbers in both seasons became more marked after the severe 1993 drought. Second, the overall average number of zebra in the Crater appeared stable whereas numbers of the other eight species declined substantially between 1974 and 2012 relative to their peak numbers during 1974-1976. Third, numbers of both gazelles, eland, kongoni, waterbuck (wet season only) and black rhino declined significantly in the Crater in both seasons following the removal of the Maasai and their cattle from the Crater in 1974. The decline in black rhino is mainly attributed to poaching in the 1970’s and 1980’s which reduced the population to 10 individuals [65]. Fourth, wildebeest numbers decreased in the Crater between 1974 and 2012 but this decrease was not statistically significant. In addition, some herbivore species were consistently more abundant inside the Crater during the wet than the dry season. This pattern was most evident for the large herbivore species requiring bulk forage, comprising buffalo, eland, elephant and black rhino. The latter may spend less time in the swamps and the forest during the wet season and may be easier to count.

### Herbivore biomass

Despite the significant changes in the population sizes of individual species in the Crater, the total herbivore biomass remained relatively stable from 1963 to 1974 and from 1974-2012, implying that the Crater has a stable multi-herbivore community. There is a tendency towards a higher biomass during the wet season, but it is not significant. Total wild herbivore biomass has not been significantly affected by the removal of the pastoralists and their livestock. The change in the grassland structure from mainly short grasses to mid to tall grasses after the removal of the Maasai and their livestock may have enhanced the forage availability for Cape buffalo, a large-bodied ruminant. The biomass of buffalo had the most dramatic increase post 1974 to become a major constituent of the total large herbivore biomass after the elimination of cattle from the Crater in 1974. A similar increase in buffalo numbers at the expense of small and medium herbivores has also been documented for Nairobi and Lake Nakuru National Parks in Kenya [66,67].

### Relationship between herbivore population size and rainfall

Rainfall significantly influenced herbivore abundance in Ngorongoro Crater and this influence varied with species and season and partly reflect functional distinctions between the species based on their life-history traits (body size, gut morphology) or life-history strategies (feeding and foraging styles). Herbivores responded to rainfall variation in three different ways in both seasons. In the wet season, numbers of herbivore species either decreased (wildebeest, eland, kongoni, waterbuck and rhino), increased (zebra, buffalo, ostrich and elephant) or increased up to intermediate levels of rainfall and then decreased with further increase in rainfall (both gazelles and warthog). Similarly, in the dry season the numbers of the herbivore species either decreased (both gazelles and waterbuck), increased (wildebeest, zebra, buffalo, ostrich and warthog) or increased up to intermediate levels of rainfall and then decreased with further increase in rainfall (eland, kongoni, elephant and rhino).

### Forecasted herbivore population dynamics

The projected population trends suggest strong interspecific contrasts regarding the scenario under which each species will likely perform best but broad similarities exist between seasons for each scenario. Except for buffalo whose numbers appear to approach asymptotes, population trajectories for wildebeest, zebra and both gazelles exhibit pronounced and sustained oscillatory dynamics, reflecting rainfall oscillations. The projected population trajectories for buffalo and zebra suggest that both species will be most abundant in the Crater under the RCP2.6 scenario, intermediate under RCP4.5 and least abundant under RCP8.5 in both seasons. This is expected since buffalo is a large-sized bulk grazer and zebra is a large-sized non-ruminant able to process large quantities of low quality forage expected to be most abundant under the wetter and cooler conditions anticipated under RCP2.6 relative to RCP4.5 and 8.5. Moreover, for both buffalo and zebra, the projected trajectories are generally similar between the RCP4.5 and 8.5 scenarios for both seasons.

By contrast, the wildebeest that requires short, green grass is anticipated to be more abundant under the RCP4.5 and 8.5 scenarios than under the RCP2.6 scenario with wetter conditions in the wet season. In the more arid dry season conditions, wildebeest should however thrive better under the more moist RCP2.6 scenario than under RCPs 4.5 and 8.5.

Trajectories for both gazelles suggest that both species will be most abundant under RCP8.5 with the lowest average rainfall, intermediate under RCP4.5 with intermediate rainfall and least abundant under RCP2.6 with the highest rainfall. This is consistent with the preference of both species for high-quality, short grasses and forbs. For both gazelles numbers will likely increase from about 2050-2060 to 2100 under RCP4.5. Also, for both gazelles, the projections suggest persistent and similar population oscillations between both seasons under each of the three scenarios. The oscillations suggest extended periods of population decline followed by increase for both gazelles in both seasons. We reiterate that these projections are based solely on rainfall influences on large herbivore population dynamics, yet the dynamics of large herbivores are often influenced by a multitude of other factors.

### Predation

The major predators in Ngorongoro Crater are lions and spotted hyenas. These species, their population dynamics and feeding ecology have been studied since the 1960’s [4-8, 68-70].

In the 1960’s the Ngorongoro Crater had a population of approximately 298 spotted hyenas [4]. When Höner et al [70] started their research in 1996 the population was about 117 hyenas and the recruitment rate was higher and the mortality was lower than during Kruuk’s study period in the 1960’s. Herbivore census data indicates that there had been a decline in the spotted hyena prey populations, i.e. wildebeest, zebra, Thomson’s gazelle, and Grant’s gazelle by 1996. From 1996 to 2002, there was an increase in the hyena population to 333 individuals. From 1996 to 2002 there was an increase in the abundance of these prey species with an average prey density of 139 ± 76 prey animals per km^2^. Höner et al [70] attribute the decline in the hyena population from the 1960’s to 1996 to the decline in their prey populations. However, from 1996 to 2002, the major predictor for the spotted hyena population increase was the increase in their prey population. Subsequently there was a reduction and then recovery of the population during an outbreak of *Streptococcus equi ruminatorum* in 2001 to 2003. Mortality was higher in adult males and yearlings in territories where prey densities were low. In the short term the bacterial infection had a top-down impact on sex and age classes that had relatively poor nutrition. In the longer-term after the disease perturbation, the reduced population growth was due to lower juvenile survival. By 2008 the population had recovered and was approximately 450 [71] and in 2012 the population was estimated at 508 of which 364 were adults (pers com Höner 2018).

From 1970 to 1972, Elliot and McTaggart Cowan [68] studied lions in the Crater and estimated a resident population of 65 lions in four prides. They estimated that lions annually killed or scavenged approximately 7% of the wildebeest, 4.3% of the zebra and 6.2% of the Thomson’s gazelle. Adapted from Kruuk [4] they estimated that hyenas took at least 7.6% of the wildebeest population, 6.5% of the zebra population and 1.6% of the Thomson’s gazelle population. Thus the estimated annual percentage of wildebeest killed or scavenged by lions and hyena was approximately 14.6%, roughly equal to the wildebeest recruitment rate [4]. The predation and scavenging rate on zebra was approximately 10.7%.

Long term research on lions in the Ngorongoro Crater [7,8,69]() indicates that the lion population may not be food limited but that weather extremes (high rainfall/drought) correlate with disease outbreaks and pest infestations (Canine distemper virus and biting *Stomoxys* flys). The resulting mortality is exacerbated by pride takeovers and infanticide. A severe infestation of *Stomoxys* flys in 1962 reduced the lion population to 10 lions that were joined by seven immigrating males in 1975. This severe population reduction may have been a ‘bottleneck’ and the current population may be based on 15 founders [7]. The population rose to a high of 124 lions in 1983, but by 1991 there were 75 to 100 lions, and numbers dropped to 29 in 1998 [7,8]. The lion population may be density dependent since it has had positive reproductive performance when the population has been less than 60 individuals and has had negative reproductive performance when the population was more than 60 individuals. From 1994 to 2004, the population had not had reduced reproductive performance. Kissui and Packer [8] attribute the declines in the lion population to disease outbreaks that correlated with extreme weather events that occurred in 1962, 1994, 1997, and 2001. During 2000/2001 there was a decrease in the lion population due to death (*Stomoxys* flys) and emigration [70].

### Poaching

The black rhino declining trend from the 1970’s to mid 1980’s was due to poaching [72]. Since the early 1990’s there has been limited poaching and the population is slowly recovering. Conservative population projections in 1995 [65] predicted that with the best scenario, i.e. no poaching, the population should be approximately 35 to 40 individuals by 2017. The current Black rhino population is 59 individuals (Pers comm, M. Musuha, 2018, NCAA,).

### Disease

Before the 1960’s, rinderpest was a source of significant mortality to buffalo, wildebeest and eland in the Crater Highlands and there was a serious outbreak affecting yearling buffalo adjacent to the Ngorongoro Crater in 1961 [73]. The NCAA started an inoculation campaign against rinderpest in the 1950’s and eradicated the disease by the 1960’s [73]. Inoculations against rinderpest for cattle continued. Subsequently there was an outbreak in 1982 that affected buffalo, eland and giraffe, but not cattle [32]. Despite the losses from rinderpest during 1982, the buffalo population increased steadily from 1980 and had doubled by 1987.

Rinderpest was also a significant source of mortality in the adjacent Serengeti ecosystem and the inoculation campaigns appear to have reduced mortality in both wildebeest populations. From 1963 to 1974 the Serengeti migratory wildebeest population tripled in size [74]. During the same time the more sedentary population in Ngorongoro Crater increased from roughly 7,600 wildebeest to about 14,000.

In 2000 and 2001 there was significant mortality in buffalo (1500), wildebeest (250) and zebra (100) apparently due to nutritional stress resulting from the severe drought in the dry season in 2000 [2, 39,75].

In 2001, five black rhinos died in January and five lions during February [62]. The reports indicated that three of the black rhinos died from *Babesiosis* [75]. Nijhof et al [75] analysed samples from Ngorongoro (Bahati and Maggie) and a dead black rhino (Benji) from Addo Elephant National Park. Sequence analyses of the sample from Bahati’s brain revealed a novel species that was named *Babesia bicornis* ap.nov. Subsequent analyses showed that both Maggie and Benji were positive for *Babesia bicornis* ap.nov. Hence, *Babesia bicornis* ap.nov. may be a species new to the Ngorongoro Crater and the Serengeti ecosystem. Two black rhino were translocated from Addo Elephant National Park to Ngorongoro Crater in 1997. The translocation that was done to enhance the Ngorongoro black rhino population may have had negative repercussions by introducing a new tick borne disease. The impact of the novel parasite, *Babesia bicornis* ap.nov., may have been exacerbated by drought and high tick densities. The literature indicates that *Babesia bicornis* can cause fatal babesiosis [75].The remaining 10 black rhinos were treated with a curative babesicidal drug and survived [39].

However, in the case of the buffalo mortalities, high tick burdens and tick borne protozoal diseases may have been contributing factors [75]. A limited survey of the buffalo, wildebeest and lions that died in 2001 did not reveal the presence of *Babesia bicornis* ap.nov. Lion necropsy’s revealed the presence of tick borne parasites (*Ehrlichia spp*., *Babesia* and *Theileria sp*) but canine distemper and a plague of *stomoxys* stinging flies were also implicated and the cause of mortality has not been determined [39].

Prescribed burning was started in the dry season of 2001 and research was done on tick densities, vegetation structure and tick host preference in adjacent burned and unburned areas [39]. Before burning, most adult ticks were present in the wet season (May to June) and most immature ticks occurred during the dry season (September, October). There were significantly more adult ticks in the tall grass in the wet season and significantly more immature ticks in the less grazed areas in the dry season. In 2001, the mean tick density in tall grass (>50cm) was 57 ± 6.93/m^2^ (adults, wet season) and in less grazed (>20 cm) areas 961 ± 146 /m^2^ (immature, dry season). Twenty-seven months (2004) later there was a significant difference between burned and unburned areas, with almost no adult ticks and relatively few immature ticks in the burned areas. However, the unburned areas also had much lower adult tick and immature tick densities than that recorded in 2001.

## Conclusions

Ngorongoro Crater has an annual rainfall cycle period of about 5 years. Oscillations in annual, wet and dry season rainfall were statistically significant. The oscillations are associated with recurrent severe droughts that cause food scarcity and hence nutritional stress for the large herbivores. Rainfall oscillations imply that large herbivores are exposed to above average food supply for about 2.5 years and to below average food supply for the subsequent 2.5 years. High rainfall supports above-average production of plant biomass which may be of low quality due to the dilution of plant nutrients.

In 1974 there was a perturbation in that resident Maasai and their livestock were removed from the Crater. Vegetation maps from before and after the removal of pastoralists and their livestock indicate that major changes in vegetation structure occurred. The 1995 vegetation map shows that there was a significant change in the vegetation structure of the Crater floor, such that there was a decrease in the availability of short grasses and an increase in medium and tall grassland.

Temporal variation in herbivore numbers in the Crater followed four general patterns. First, buffalo, elephant and ostrich numbers increased significantly in the Crater from 1974-2012. Second, the overall average number of zebra in the Crater appeared stable whereas numbers of the other eight species declined substantially between 1974 and 2012 relative to their peak numbers during 1974-1976. Third, numbers of both gazelles, eland, kongoni, waterbuck (wet season only) and black rhino declined significantly in the Crater in both seasons following the removal of the Maasai and their cattle from the Crater in 1974. The decline in black rhino is mainly attributed to poaching in the 1970’s and 1980’s. Fourth, wildebeest numbers decreased in the Crater between 1974 and 2012 but this decrease was not statistically significant. In addition, some herbivore species were consistently more abundant inside the Crater during the wet than the dry season. This pattern was most evident for the large herbivore species requiring bulk forage, comprising buffalo, eland, elephant and black rhino. The latter may spend less time in the swamps and the forest during the wet season and may be easier to count. Even with a change in grassland structure, total herbivore biomass remained relatively stable from 1963 to 2012, implying that the Crater has a stable multi-herbivore community.

Rainfall significantly influenced herbivore abundance in Ngorongoro Crater and this influence varied with species and season. Herbivores responded to rainfall variation in three different ways in both seasons. In the wet season, numbers of herbivore species either decreased (wildebeest, eland, kongoni, waterbuck and rhino), increased (zebra, buffalo, ostrich and elephant) or increased up to intermediate levels of rainfall and then decreased with further increase in rainfall (both gazelles and warthog). Similarly, in the dry season the numbers of the herbivore species either decreased (both gazelles and waterbuck), increased (wildebeest, zebra, buffalo, ostrich and warthog) or increased up to intermediate levels of rainfall and then decreased with further increase in rainfall (eland, kongoni, elephant and rhino).

The relationships established between the time series of historic animal counts in the wet and dry seasons and lagged wet and dry season rainfall series were used to forecast the likely future trajectories of the wet and dry season population size for each species under three alternative climate change scenarios. They suggest strong interspecific contrasts regarding the scenario under which each species will likely perform best but broad similarities exist between seasons for each scenario.

There is information on the population trends of the two major predators, i.e. lions and spotted hyenas. It would be useful to correlate predator impact on herbivore populations with rainfall. Disease is an important perturbation in the population trends of lions and spotted hyenas and potentially Black rhino, Cape buffalo and other herbivores. Tick borne diseases can potentially be managed with systematic burning of some grassland areas.

## Acknowledgements

JOO was supported by a grant from the German National Research Foundation (Grant No. OG 83/1-1). JOO was also supported The Planning for Resilience in East Africa through Policy, Adaptation, Research, and Economic Development (USAID PREPARED) project. This project has received funding from the European Union’s Horizon 2020 research and innovation programme under Grant Agreement No. 641918. PDM is grateful for permission from the Tanzania Commission for Science and Technology (COSTECH) and the Tanzania Wildlife Research Institute (TAWIRI) to conduct long-term research in the Serengeti–Ngorongoro ecosystem. TAWIRI, the Tanzania National Parks Authority (TANAPA) and the Ngorongoro Conservation Area Authority (NCAA) have provided important support for this research.

## Supporting Information

S1 Data. The count totals for each of the 12 most common large herbivore species counted during the wet and the dry seasons in the Ngorongoro Crater from 1964 to 2012.

**S2 Data. The count totals for each of the 12 most common large herbivore species counted during the wet and the dry seasons in the Ngorongoro Crater from 1964 to 2012**. The missing values were imputed using a state space model, separately for each species and season combination.

S3 Data. Total monthly rainfall in mm recorded at the Ngorongoro Conservation Area headquarters from 1963 to 2014.

**S4 Data. The logarithm of the observed and predicted population size for each of the five most common species for the wet and dry season and the 95% pointwise prediction confidence band for 1964 to 2012**. The logarithm of the forecasted population size is also provided for each of the five most abundant herbivore species for 2013 to 2100.

**Table S1. Parameter estimates for the bivariate VARMAX (2,2,5) model for the five most abundant herbivore species in the dry and wet seasons in the Ngorongoro Crater, Tanzania, during 1963-2012**. Model selection was based on information theory so no effort has been made to remove insignificant coefficients. By restricting a few of the highly insignificant coefficients to be zero, many of the apparently insignificant coefficients become significant.

**Table S2. Roots of AR characteristic polynomials for the bivariate model for the five most abundant herbivore species in the dry and wet seasons in the Ngorongoro Crater, Tanzania, during 1963-2012**. The modulus of the roots of its AR polynomial should be less than 1 for a time series to be stationary.

Table S3. Roots of the MA characteristic polynomials for the bivariate model for the five most abundant herbivore species in the dry and wet seasons in the Ngorongoro Crater, Tanzania, during 1963-2012.

**Table S4. Portmanteau Test for Cross Correlations of Residuals from the bivariate VARMAX(2,2,5) model for the five most abundant herbivore species in the dry and wet seasons in the Ngorongoro Crater, Tanzania, during 1963-2012**. The results show tests for white noise residuals based on the cross correlations of the residuals. Insignificant test results show that we cannot reject the null hypothesis that the residuals are uncorrelated.

**Table S5. Univariate model ANOVA diagnostics for the five most abundant herbivore species in the dry and wet seasons in the Ngorongoro Crater, Tanzania, during 1963-2012**. The results show that each model is significant.

**Table S6. Univariate Model White Noise Diagnostics for the five most abundant herbivore species in the dry and wet seasons in the Ngorongoro Crater, Tanzania, during 1963-2012**. The results show tests of whether the residuals are correlated and heteroscedastic. The Durbin-Watson test statistics test the null hypothesis that the residuals are uncorrelated. The Jarque-Bera normality test tests the null hypothesis that the residuals are normally distributed. The F statistics and their *p*-values for ARCH(1) disturbances test the null hypothesis that the residuals have equal covariances.

**Table S7. Univariate AR Model Diagnostics for the five most abundant herbivore species in the dry and wet seasons in the Ngorongoro Crater, Tanzania, during 1963-2012**. The F statistics and their *p*-values for AR(1), AR(1,2), AR(1,2,3) and AR(1,2,3,4) models of residuals test the null hypothesis that the residuals are uncorrelated.

Table S8. Classification of years and seasons into extreme drought, severe drought, moderate drought, normal, wet, very wet and extremely wet years or seasons using percentiles of the frequency distributions of the total annual, wet season or dry season rainfall recorded at the Ngorongoro Conservation Area headquarters from 1963 to 2014.

Table S9. The estimated frequency, period, periodogram, spectral density, co-spectra, quadrature, squared coherence, amplitude and phases of the oscillations in the annual, wet and dry season rainfall components for the Ngorongoro Crater during 1963-2014.

Table S10. The estimated variances of the disturbance terms, the variances of the irregular components, damping factor and periods of the cycles in the annual, wet and dry season rainfall components recorded for the Ngorongoro Crater during 1963-2014.

Table S11. Significance analysis of components (based on the final state).

Table S12. The expected population size of each of the 12 wildlife species in 1964, 1974 and 2012 and the difference between the two estimates for 1964 and 1974 and 1974 and 2012 and test of significance of their difference based on constructed penalized cubic B-splines.

Table S13. Selection of the rainfall component, moving average and functional form of the relationship between population size and the moving average component for each of the 12 most common large herbivore species based on the corrected Akaike Information Criterion (AICc). Only models with delta AICc no more than 4 are shown. Model selection was carried out separately for the wet and dry season counts for each species.

Table S14. Parameters estimates, their standard errors and *t*-tests of whether the parameters are significantly different from zero for the AICc-selected best models relating population size and moving average rainfall, for the wet and season counts, for the 12 most common large herbivore species in the Ngorongoro Crater.

S1 Text. SAS code used to analyze the rainfall data for the Ngorongoro Conservation Area headquarters.

S2 Text. SAS code used to model trends in the animal counts, relate the counts to rainfall and project population dynamics to 2013-2100.

Fig S1. Temporal variation in the original and smoothed total monthly rainfall in the Ngorongoro Crater from 1963 to 2014.

Fig S2. Percentiles of the annual, dry and wet season rainfall components. The percentiles are used to classify years or seasons as extreme, severe or moderate drought years or seasons, normal, wet, very wet or extremely wet years or seasons as described in the text.

**Fig S3. Spectral density versus period of cycles (in years) for a) annual rainfall, b) wet season rainfall, and c) dry season rainfall based on rainfall recorded for the Ngorongoro Conservation Authority headquaters from 1963 to 2014**. A large value of spectral density means that the corresponding period has greater support in the data.

Fig S4. Smoothed cycles and trends based on the structural time series analysis versus the year of observation for the standardized annual (annualstd), wet season (wetstd) and dry season (drystd) rainfall for the Ngorongoro Crater for 1963-2014.

Fig S5. Projected total annual rainfall, average maximum and minimum temperatures for Ngorongoro Crater in Tanzania under three climate scenarios (RCP2.6, RCP4.5 and RCP8.5) for the period 2006-2100.

